# Examining the composition of gut microbiota in a South African population: a comparative study between type 2 diabetes mellitus patients and non-diabetic individuals

**DOI:** 10.1101/2025.07.16.665184

**Authors:** Sara Mosima Pheeha, Justice T. Ngom, Abhinav Sharma, Bettina Chale-Matsau, Kristien Nel Van Zyl, Samuel Manda, Peter Suwirakwenda Nyasulu

## Abstract

**Background:** Literature has highlighted the gut microbiota’s role in metabolic functions, suggesting a potential link between gut microbiota composition and T2DM. The purpose of the study was to identify microbial signatures unique to T2DM patients and non-diabetic individuals, to compare microbial profiles between the two groups and to investigate how gut microbiota may be related to inflammation associated with T2DM.

**Methods:** A cross-sectional study was conducted involving 51 T2DM patients and 99 non-diabetic South African individuals. Faecal samples were collected and analysed using 16S rRNA gene sequencing to characterize the gut microbiota. Blood samples were obtained to perform HbA1c, CRP and ferritin tests. Bioinformatic and statistical analyses were performed to identify differences in microbial composition and diversity between the two groups.

**Results:** The gut microbiota in T2DM patients was predominantly composed of Firmicutes (47.7%), Bacteroidota (37.5%), and Proteobacteria (11.4%), while the non-diabetic group showed a slightly different microbial profile with higher Bacteroidota (41.9%) and a notable presence of Actinobacteriota (4.5%). Abundant families in the T2DM group included Bacteroidaceae (22.8%), Prevotellaceae (7.4%), Enterobacteriaceae (7.4%), Erysipelotrichaceae (6.0%) and Lachnospiraceae (5.2%). The non-diabetic group exhibited dominant families such as Lachnospiraceae 26.7%, Prevotellaceae (25.3%), Bacteroidaceae (12.7%), Ruminococcaceae (9.5%) and Oscillospiraceae (3.8%). At the genus level, Bacteroides (22.8%), Escherichia-Shigella (5.0%), Holdemanella (4.8%), Phascolarctobacterium (3.2%) and Blautia (2.8%) were prevalent in the T2DM group, while Prevotella_9 (22.1%), Bacteroides (12.7%), Agathobacter (6.7%), Blautia (6.3%) and Faecalibacterium (5.1%) were dominant in the non-diabetic group. Differential abundance testing revealed 5 phyla, 16 families, and 25 genera that were either enriched/depleted in T2DM patients relative to non-diabetic individuals. The comparison of alpha diversity metrics between the two groups revealed significant differences across all four measures (P < 0.001), with non-diabetic individuals showing higher values than T2DM patients. HbA1c and CRP levels showed correlations with the relative abundance of various gut microbes at various phyla, family, and genus levels, as well as with all alpha diversity metrics.

**Conclusion:** The study revealed distinct differences in gut microbiota composition between T2DM patients and non-diabetic individuals, with T2DM patients showing a higher prevalence of certain phyla, families, and genera linked to metabolic dysregulation. Non-diabetic individuals exhibited greater microbial diversity and beneficial taxa, highlighting a potential protective microbial profile.

## 1. Introduction

The human gastrointestinal tract is an intricate ecosystem inhabited by trillions of microorganisms collectively known as the gut microbiota [1]. The complex relationship between human health and the gut microbiota has attracted remarkable attention in recent years, as emerging research explains the important role these microbial communities play in metabolic and immune processes [2]. Among the numerous health conditions influenced by the gut microbiota, type 2 diabetes mellitus (T2DM) stands as an alarming global health challenge [3], affecting millions of individuals and posing a substantial burden on healthcare systems worldwide [4].

The escalating prevalence of T2DM, particularly in regions undergoing rapid epidemiological transitions or urbanization, changing dietary patterns and genetic diversity, like South Africa [5], necessitates a thorough understanding of the underlying factors contributing to its emergence and progression. Gaining insights into the core factors that contribute to T2DM in this population is essential for the development of inexpensive as well as effective prevention and management strategies.

Recent research has increasingly focused on the potential link between gut microbiota composition and metabolic disorders, including T2DM [6]. Several studies have reported on alterations in gut microbial profiles among patients with T2DM when compared to their non-diabetic counterparts [7–10]. However, despite a growing body of research explaining the association between gut microbiota and T2DM, there exists a lack of studies conducted in the African continent [10,11], specifically related to the unique demographic and environmental context of South Africa. Investigating gut microbiota across diverse populations is crucial due to variations in ethnicity and lifestyle practices. Ethnic and lifestyle differences may contribute to unique patterns in gut microbiota, and exploring these variations can provide valuable insights into the relationship between gut microbiota and T2DM.

This study aimed to address this critical gap by conducting a comprehensive comparative analysis of gut microbiota composition in a South African population, examining both patients diagnosed with T2DM and their non-diabetic counterparts. By investigating the specific microbial signatures associated with T2DM in this distinct demographic, we aspire to discover potential microbial changes that could be pivotal in understanding the pathophysiology of T2DM within the South African setting.

Furthermore, this study sought to shed light on the potential correlations between inflammatory markers (C-reactive protein, CRP and Ferritin) and gut microbiota profiles in this population. Mounting evidence suggests that alterations in the composition and diversity of gut microbiota, known as dysbiosis, can have a remarkable impact on the development and progression of T2DM [12]. Dysbiosis can lead to increased intestinal permeability and the translocation of bacterial products into the systemic circulation [13]. These products (i.e., lipopolysaccharides) [13] activate toll-like receptors triggering a cascade of inflammatory responses [14]. Consequently, elevated levels of pro-inflammatory cytokines such as interleukin-6 (IL-6), tumour necrosis factor-alpha (TNF-α), and CRP have been observed in patients with T2DM [15]. Moreover, this chronic inflammation is closely associated with insulin resistance [16], impaired glucose metabolism [17] and pancreatic β-cell dysfunction [18].

Additionally, some types of beneficial microorganisms (e.g. *Faecalibacterium prausnitzii*, *Eubacterium rectale* and *Eubacterium hallii* [19]) in the gut generate distinct enzymes that facilitate the transformation of nutrients into absorbable forms, such as converting indigestible carbohydrates into short chain fatty acids (SCFAs), which might exhibit properties with anti-inflammatory and immunomodulatory effects [20]. Patients diagnosed with T2DM often display a higher presence of harmful gut bacteria relative to beneficial ones. This imbalance can lead to reduced production of SCFAs, which are acknowledged for their anti-inflammatory attributes [21]. Understanding the dynamic relationship between gut microbiota and inflammatory markers in T2DM provides crucial insights into its pathophysiology. It also holds promise for the development of targeted interventions aimed at modulating the gut microbiome to mitigate inflammation and improve metabolic outcomes in patients with this prevalent metabolic disorder.

To the best of our knowledge, this study is the first of its kind in South Africa, and the findings may have the potential to contribute significantly to the global understanding of the role of gut microbiota in metabolic disorders and offer crucial insights into the specific microbial determinants of T2DM within the South African population. These insights may pave the way for targeted interventions and personalized approaches for the management of T2DM in this population.

## 2. Materials and Methods

### Sample collection

Blood and stool samples were obtained from all the participants. Samples from patients with diabetes (*n*= 51) were collected at the diabetic clinic of DGMAH. Samples from non-diabetic individuals (*n*= 99) were collected at the Sefako Makgatho Health Sciences University’s Chemical Pathology Department. Although the samples were collected in batches, the same sample collection protocol was adhered to for all participants in the study. Blood was drawn using EDTA and BD Vacutainer™ venous blood collection tubes, specifically SST™ Serum Separation tubes, in both groups. Stool samples were obtained using OMNIgene•GUT OM- 200 kits (DNA Genotek Inc., Canada), as per the manufacturer’s instructions. Specimens did not have to be frozen after collection and while being transported to Inqaba Biotechnical Industries (Pty) Ltd laboratory for 16S rRNA gene sequencing analysis. This is because the collection kits included a solution with a chelating agent to effectively stabilise DNA present in faecal samples, making collection, prolonged storage, and transportation at ambient temperature possible.

Diabetic patients who could not provide stool samples at their clinic visit were supplied with faecal collection kits and guided to collect them at their convenience in their homes. The researchers used the provided home addresses to trace the patient and collected the samples once the patients signalled by phone call that they had collected the stool sample. Every non-diabetic participant managed to provide stool samples promptly as per the recruitment instruction to visit the department only when they were prepared to pass stools.

### Data collection and biochemical tests

We collected demographic data including age, gender, location, diabetes mellitus status, physical activity, diet, type of diabetes medical treatment, antibiotic use, probiotic use, and presence of gastrointestinal disorders. Additionally, participants’ whole blood samples were assessed for glycated haemoglobin (HbA1c), while serum samples were used to evaluate Ferritin and CRP levels. All biochemical assays were performed using the Atellica Solutions autoanalyser (Siemens Healthineers, Germany).

### Bacterial genomic DNA extraction

Genomic DNA was extracted using ZymoBIOMICS™ DNA Miniprep kit (Zymo Research, USA) following the manufacturer’s protocol. To determine the purity and estimate the concentration of the extracted DNA, the NanoDrop Spectrophotometer (NanoDrop Technologies, USA) was used. The resulting amplicons were also checked for quality on agarose gel electrophoresis and quantified with Qubit® dsDNA HS (High Sensitivity) Assay kits.

### Library preparation for amplicon sequencing

Polymerase chain reaction was performed using universal primer pair 27F and 1492R, targeting the V1 -V9 region of the bacterial 16S rRNA gene. These primers were tagged with PacBio M13 adaptor sequences on the 5’-end of the forward and reverse sequences to allow barcoding of each amplicon. Resulting amplicons were barcoded with PacBio M13 barcodes for multiplexing through limited cycle PCR. The resulting barcoded amplicons were quantified and pooled equimolar, and the AMPure PB bead-based purification step was performed.

### PacBio SMRTbell library preparation, metagenomic analysis of full length 16S gene amplicons and bioinformatics analysis

Samples were sequenced on the Sequel II system by PacBio (www.pacb.com). Raw sub-reads were processed through the SMRTlink (v11.0) Circular Consensus Sequences (CCS) algorithm to produce highly accurate reads (>QV40). These reads were then analysed in a reproducible manner using the Nextflow [22] based nf-core community [23] pipeline ampliseq (v2.6.1, https://nf-co.re/ampliseq/2.6.1) [24] with both DADA2 (https://benjjneb.github.io/dada2/index.html) and QIIME2 v2023.7 (https://qiime2.org/) options enabled for quality control assessment, taxonomic classification as well as relative abundance analysis. Clustering, denoising and filtering was performed using DADA2 with the default parameters and truncating reads at mean quality < Q10. Cutadapt was used to remove primers sequences. The SILVA 138.1 full-length prokaryotic SSU database was used as reference for taxonomic classification. The approach also enabled the assessment of microbial community diversity (alpha and beta diversity metrics). Alpha diversity was determined using the Shannon’s diversity index, Faith’s PD, Pielou’s evenness index and observed features. Beta diversity (microbial community dissimilarity) investigation was conducted with principal coordinate analysis (PCoA) based on the Bray-Curtis metric, weighted and unweighted UniFrac distance. Permutational multivariate analysis of variance (PERMANOVA) was conducted using distance matrices. Adonis was employed to assess the significant differences between the group with T2DM and the non-diabetic group. Furthermore, differential abundance testing was performed using the Analysis of Compositions of Microbiomes with Bias Correction (ANCOM-BC) plugin in QIIME2 v2023.7. The versions of individual software and exact parameters, as well as the taxonomic databases used in the pipeline are documented in the supplementary (Table S1) MultiQC version 1.18 report [25]. Finally, potential confounding associations between bacterial features and covariates were explored with MaAsLin2 at the lowest taxonomic level reported in this study (genus), which was most sensitive to change. The genus level counts produced by the pipeline described above were transformed to relative abundance, whereafter a compound Poisson linear model (CPLM) was applied with covariates as fixed effects.

### Quality and contamination control

The ZymoBIOMICS Microbial Community DNA standard (Zymo Research, USA) served as an ideal positive mock control and ensured the quality of microbiome profiling. It allowed for benchmarking and validation of microbial composition profiling workflows and facilitated the assessment of bias in DNA isolation.

### Statistical analysis

Statistical analyses were conducted using STATA statistical software. The three taxa levels (Phyla, Family and Genera) were skewed, so the Mann–Whitney U test was used to investigate differences in their abundance levels between the two groups of participants. Alpha diversity metrics (Shannon index, Faith’s PD, Pielou’s evenness index and Observed features) as well as Firmicutes/Bacteroidota (F/B) ratios were also compared using the same test. The observed significant values were adjusted for multiple testing. The ANCOM-BC plugin computed an internal statistic (W), utilized for generating P-values with two-sided z tests. Subsequently, these P-values were adjusted for multiple hypothesis correction. Differentially abundant taxa with adjusted P-values (Q-values) < 0.05 were shown in the visualizations. In addition to employing STATA and QIIME2, the visual representations in this study were also created using RStudio (RStudio Team (2020). RStudio: Integrated Development for R. RStudio, PBC, Boston, MA URL http://www.rstudio.com/.) and GraphPad Prism version 10.1.2. (specifically, for generating pie charts and heat maps). A binary logistic regression analysis was performed to assess the association between various taxa and alpha diversity metrics with both T2DM status and the type of T2DM treatment administered. We used the Spearman’s correlation test to examine the linear association of HbA1c and inflammatory markers (CRP and ferritin) with relative abundance of predominant taxa and alpha diversity metrics. A P value of < 0.05 was regarded as statistically significant.

## 3. Results

The study included a total of 150 individuals, comprising 51 T2DM and 99 non-diabetic participants. Among the T2DM group, 49% of patients were on metformin as their primary antidiabetic treatment, while 25% were on insulin therapy. Furthermore, the study included 88 female participants and 59 male participants (Table 1). However, despite this variation, no significant association was found between gender and diabetes status. Age was associated with DM status, P <0.001 (Table 1). Almost half (49%) of the diabetic group displayed poor glycaemic control (HbA1c > 8%) [26], whereas the non-diabetic group (83%) primarily had normal HbA1c levels [26] (Table 1). The diabetic cohort (44%) mostly presented with moderately elevated CRP levels [27], while the non-diabetic group (55%) mostly presented with normal CRP levels [27] (Table 1).

**Table 1.** Demographic characteristics of study participants.

| Variable |  | T2DM patients<br>(n = 51) n (%) | Non-diabetic<br>individuals<br>(n = 99) n (%) | P |
| --- | --- | --- | --- | --- |
| Gender | Male | 20 (39) | 39 (39) | 1.000 |
|  | Female | 31 (61) | 57 (58) |  |
|  | Unknown | - | 3 (3) |  |
| Age [years] |  | 57 (22) | 37.5 (24) | <0.001* |
| Residence | Township | 50 (98) | 93 (94) | 0.529 |
|  | Urban | 0 (0) | 2 (2) |  |
|  | Rural | 1 (2) | 1 (1) |  |
|  | Unknown | - | 3 (3) |  |
| Type of treatment | Metformin | 25 (49) | - | - |
|  | Insulin | 13 (25) | - |  |
|  | Metformin + insulin | 7 (14) | - |  |
|  | Unknown | 6 (12) | - |  |
| HbA1c [%] | Normal ($\leq 5.6$) | 4 (8) | 80 (83) | <0.001* |
|  | Pre-diabetes (5.7- 6.4) | 6 (12) | 13 (14) |  |
| | Diabetes ($\geq 6.5$) | 16 (31) | 1 (1) | |
| | Uncontrolled diabetes ($> 8$) | 25 (49) | 2 (2) | |
| CRP (mg/L) | Normal ($< 3$) | 11 (22) | 53 (55) | <0.001* |
|  | Minor elevation (3- 10) | 15 (30) | 29 (30) |  |
|  | Moderate elevation (10- 100) | 22 (44) | 14 (15) |  |
|  | Marked elevation (100-500) | 2 (4) | 0 (0) |  |
| | Severe elevation ($> 500$) | 0 (0) | 0 (0) | |
| Ferritin in females (ug/L) | No hyperferritinaemia ($\leq 200$) | 24 (80) | 54 (98) | 0.009* |
| | Hyperferritinaemia ($> 200$) | 6 (20) | 1 (2) | |
| Ferritin in males (ug/L) | No hyperferritinaemia ($\leq 400$) | 18 (90) | 40 (98) | 1.000 |
| | Hyperferritinaemia ($> 400$) | 2 (10) | 1 (2) | |

Age is reported as median and interquartile range (IQR) while the rest are reported as number of observations (n) and percentage (%). The asterisk (\*) denotes $P < 0.05$ . Some data points for
HbA1c, CRP, and ferritin were missing, leading to calculations being performed based on 147, 146, and 146 observations for each respective variable.

### 3.1. Sequencing results

The sequencing run produced a total of 4542036 reads and after quality control and filtering, 4224201 reads (93 %) remained for downstream analysis through DADA2, retaining between 42.06% and 94.53% reads per sample (average 85.1%).

### 3.2. Phyla composition

The analysis of gut microbiota in patients with T2DM revealed the abundance of 5 major phyla: Firmicutes (47.7%), Bacteroidota (37.5%), Proteobacteria (11.4%), Actinobacteriota (1.5%) and Fusobacteriota (1.3%) (Figure 1A and 2). In contrast to the T2DM group, the composition of gut microbiota in the non-diabetic group demonstrated a slightly distinct profile which was: Firmicutes (48.5%), Bacteroidota (41.9%), Actinobacteriota (4.5%), Proteobacteria (3.4%) and Verrucomicrobiota (1.2%) (Figure 1D and 2).

**Figure 1.**
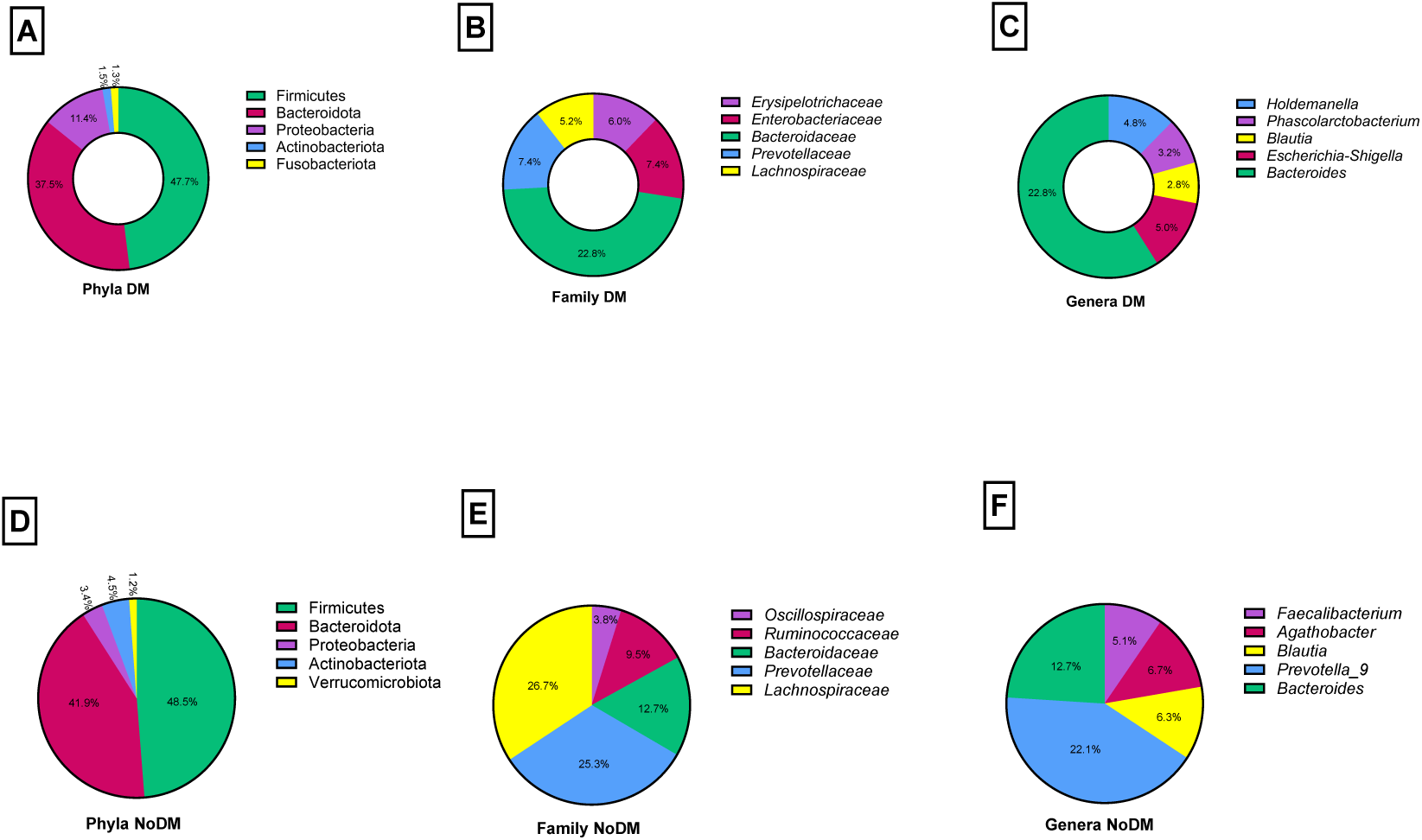
Comparison of relative abundance between T2DM patients and non-diabetic individuals at the phylum, family, and genus level. DM: diabetes mellitus: No DM: no diabetes mellitus.

**Figure 2.**
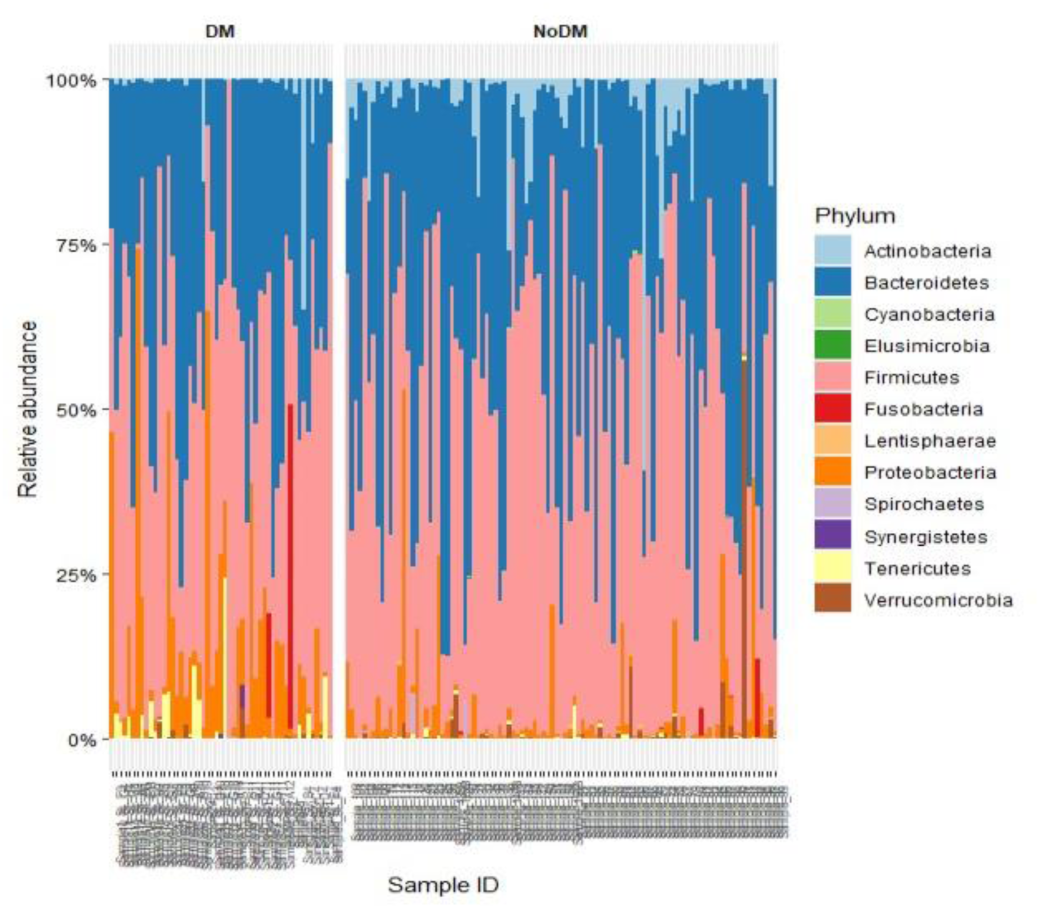
Comparison of relative abundance between T2DM patients and non-diabetic individuals at the phylum level. DM: diabetes mellitus: No DM: no diabetes mellitus.

### 3.3. Family composition

Among the T2DM group, the most abundant families were *Bacteroidaceae*, accounting for 22.8% of the microbial community, followed by *Prevotellaceae* and *Enterobacteriaceae*, each at 7.4%. Other notable families included *Erysipelotrichaceae* and *Lachnospiraceae* representing 6.0% and 5.2% of the community, respectively (Figure 1b and Figure 3). The non-diabetic group exhibited a different composition of dominant families. *Lachnospiraceae* was the most abundant, representing 26.7% of the microbial community, followed closely by *Prevotellaceae* at 25.3%. *Bacteroidaceae, Ruminococcaceae* and *Oscillospiraceae* each represented 12.7%, 9.5% and 3.8% respectively (Figure 1E and Figure 3).

**Figure 3.**
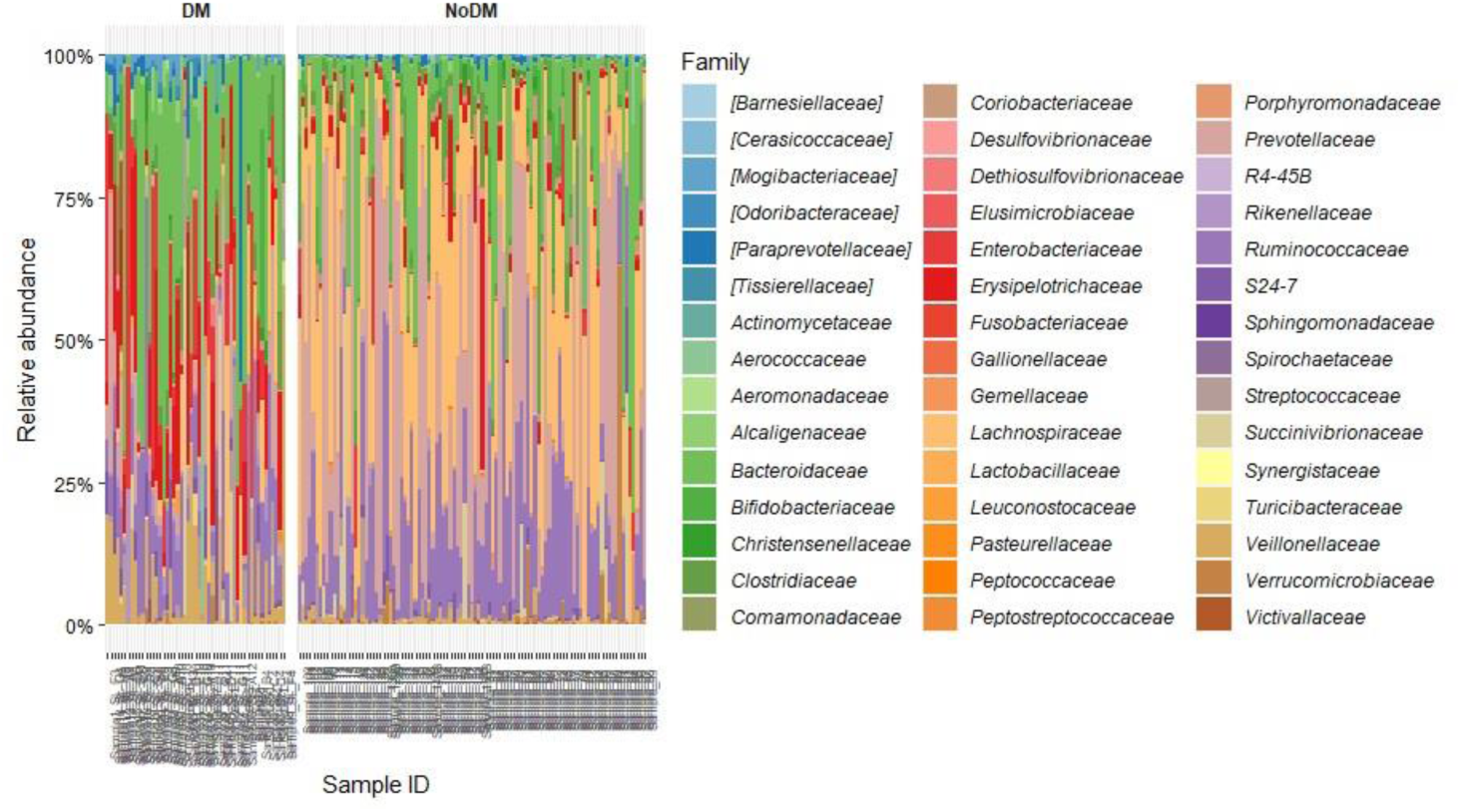
Comparison of relative abundance between T2DM patients and non-diabetic individuals at the family level. DM: diabetes mellitus: No DM: no diabetes mellitus.

### 3.4. Genera composition

The most abundant genera identified in T2DM patients included *Bacteroides*, representing 22.8% of the microbial community. *Escherichia-Shigella* constituted 5.0%, while *Holdemanella* and *Phascolarctobacterium* were both present at 4.8% and 3.2%, respectively. Additionally, *Blautia* were identified at 2.8% (Figure 1c and 4). In contrast, non-diabetic individuals exhibited a different composition of abundant genera within their gut microbiota. *Prevotella_9* was the most prevalent, representing 22.1% of the microbial community. *Bacteroides* constituted 12.7%, while *Agathobacter* accounted for 6.7%. *Blautia* was present at 6.3%, followed by *Faecalibacterium* at 5.1% (Figure 1F and 4). Table 2 presents a comparison of taxa abundance across phylum, family, and genus levels

**Figure 4.**
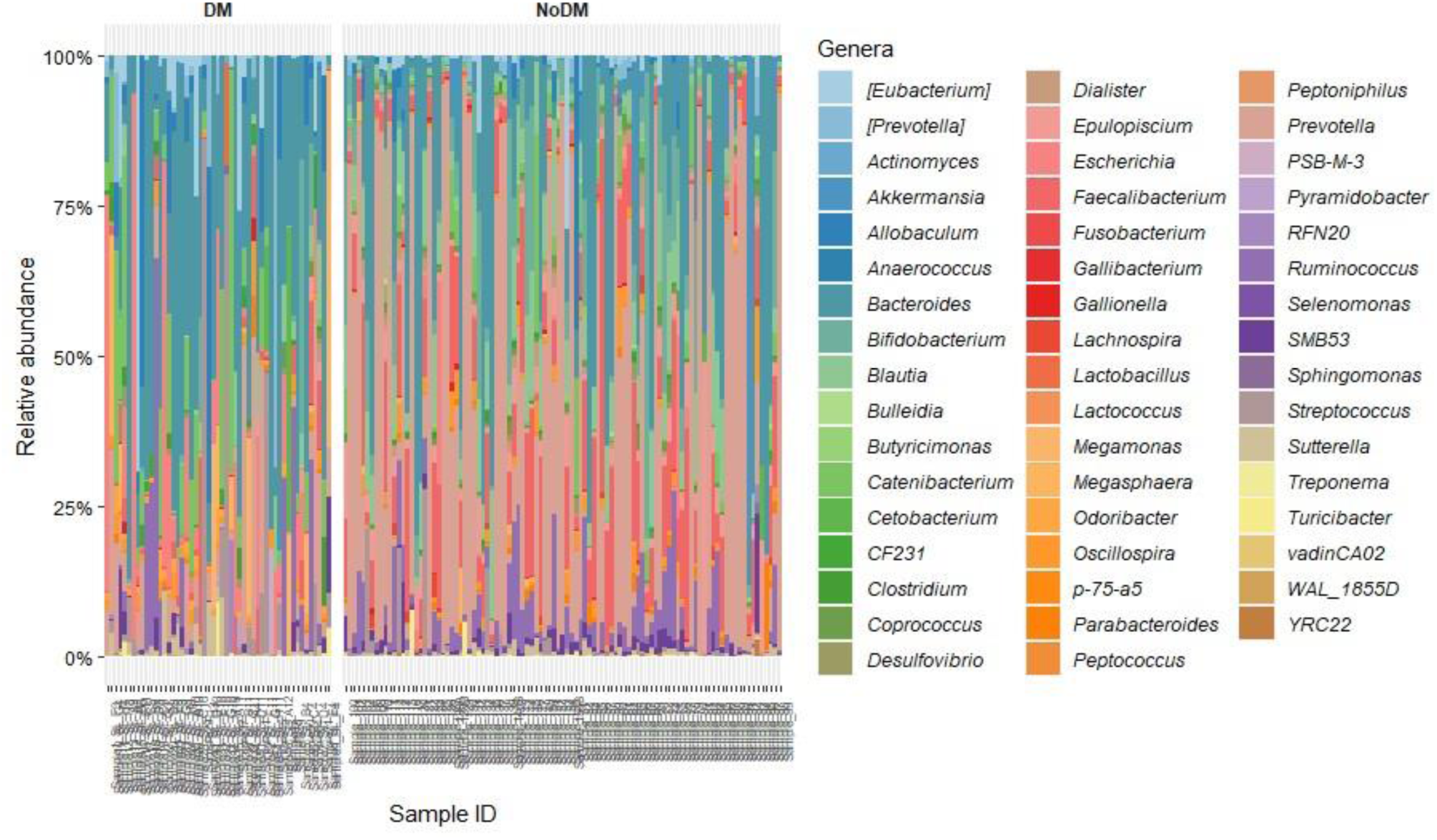
Comparison of relative abundance between T2DM patients and non-diabetic individuals at the genus level. DM: diabetes mellitus: No DM: no diabetes mellitus.

**Table 2.**
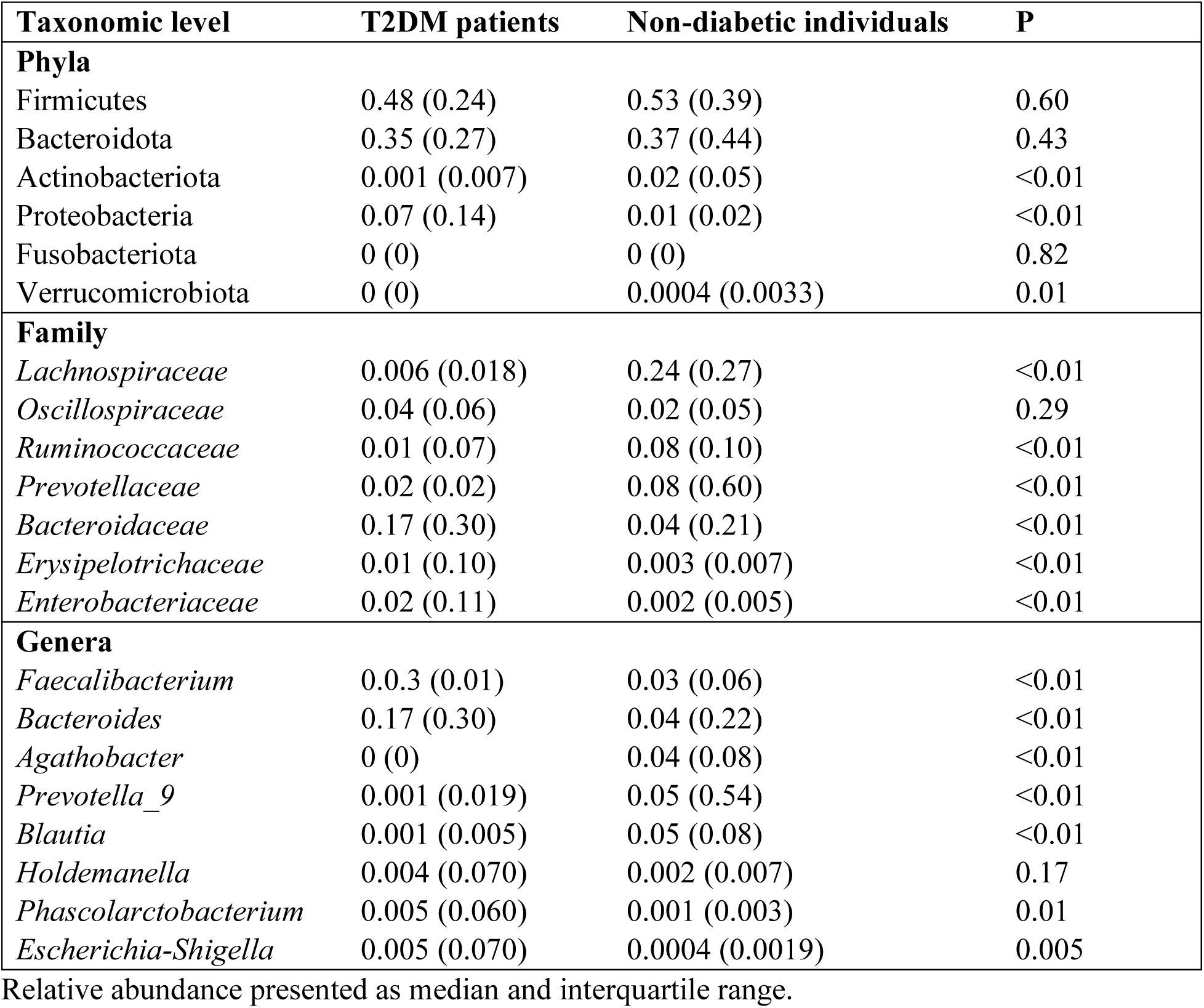
Comparison of relative abundance of taxa at phylum, family, and genus level.

| Taxonomic level | T2DM patients | Non-diabetic individuals | P |
| --- | --- | --- | --- |
| **Phyla** |  |  |  |
| Firmicutes | 0.48 (0.24) | 0.53 (0.39) | 0.60 |
| Bacteroidota | 0.35 (0.27) | 0.37 (0.44) | 0.43 |
| Actinobacteriota | 0.001 (0.007) | 0.02 (0.05) | <0.01 |
| Proteobacteria | 0.07 (0.14) | 0.01 (0.02) | <0.01 |
| Fusobacteriota | 0 (0) | 0 (0) | 0.82 |
| Verrucomicrobiota | 0 (0) | 0.0004 (0.0033) | 0.01 |
| **Family** |  |  |  |
| *Lachnospiraceae* | 0.006 (0.018) | 0.24 (0.27) | <0.01 |
| *Oscillospiraceae* | 0.04 (0.06) | 0.02 (0.05) | 0.29 |
| *Ruminococcaceae* | 0.01 (0.07) | 0.08 (0.10) | <0.01 |
| *Prevotellaceae* | 0.02 (0.02) | 0.08 (0.60) | <0.01 |
| *Bacteroidaceae* | 0.17 (0.30) | 0.04 (0.21) | <0.01 |
| *Erysipelotrichaceae* | 0.01 (0.10) | 0.003 (0.007) | <0.01 |
| *Enterobacteriaceae* | 0.02 (0.11) | 0.002 (0.005) | <0.01 |
| **Genera** |  |  |  |
| *Faecalibacterium* | 0.0.3 (0.01) | 0.03 (0.06) | <0.01 |
| *Bacteroides* | 0.17 (0.30) | 0.04 (0.22) | <0.01 |
| *Agathobacter* | 0 (0) | 0.04 (0.08) | <0.01 |
| *Prevotella_9* | 0.001 (0.019) | 0.05 (0.54) | <0.01 |
| *Blautia* | 0.001 (0.005) | 0.05 (0.08) | <0.01 |
| *Holdemanella* | 0.004 (0.070) | 0.002 (0.007) | 0.17 |
| *Phascolarctobacterium* | 0.005 (0.060) | 0.001 (0.003) | 0.01 |
| *Escherichia-Shigella* | 0.005 (0.070) | 0.0004 (0.0019) | 0.005 |

Relative abundance presented as median and interquartile range.

### 3.5. Differential abundance testing between T2DM patients and non-diabetic individuals

Differential abundance testing using the ANCOM-BC plug-in in QIIME2 was performed and 5, 16, and 25 differentially abundant features (P < 0.05) were identified at phylum, family, and genus level, respectively (Figure 5). The phyla Proteobacteria and Tenericutes were enriched in the T2DM group, and Actinobacteria, Cyanobacteria, and Lentisphaerae were depleted. Similar to what was observed using relative abundance, *Lachnospiraceae* and *Prevotellaceae* were depleted in the T2DM group, compared to non-diabetic individuals. The families *Rikenellaceae* and *Enterobacteriaceae* were observed to be significantly enriched in T2DM patients. At genus level, *Escherichia* and *Odoribacter* were enriched in the T2DM group compared to non-diabetic group, whereas *Epulopiscium* and *Blautia* were depleted. A large number of unique, but unannotated, genera were found to be differentially abundant between the study groups. Additional model-based testing to identify bacterial features associated with T2DM was performed, which also included age as a fixed effect, to assess the potential bias introduced by the age of T2DM participants, which was statistically higher than the non-diabetic participants. Of the 25 microbial features detected by ANCOM-BC, 19 (76%) were also identified as statistically significant with MaAsLin2 (Q < 0.05). In addition, eight more statistically significant features were identified, of which six were enriched in T2DM (*Butyricimonas*, Bacteroidales S24.7 genus, Clostridiales genus, *Oscillospira*, *Catenibacterium*, and *Enterobacteriaceae* genus) and two were depleted (*Barnesiellaceae* genus, *Victivallaceae* genus). Only one genus level feature associated with T2DM (*Clostridiaceae* SMB53), was also found to be enriched significantly with increased age, and was excluded.

**Figure 5.**
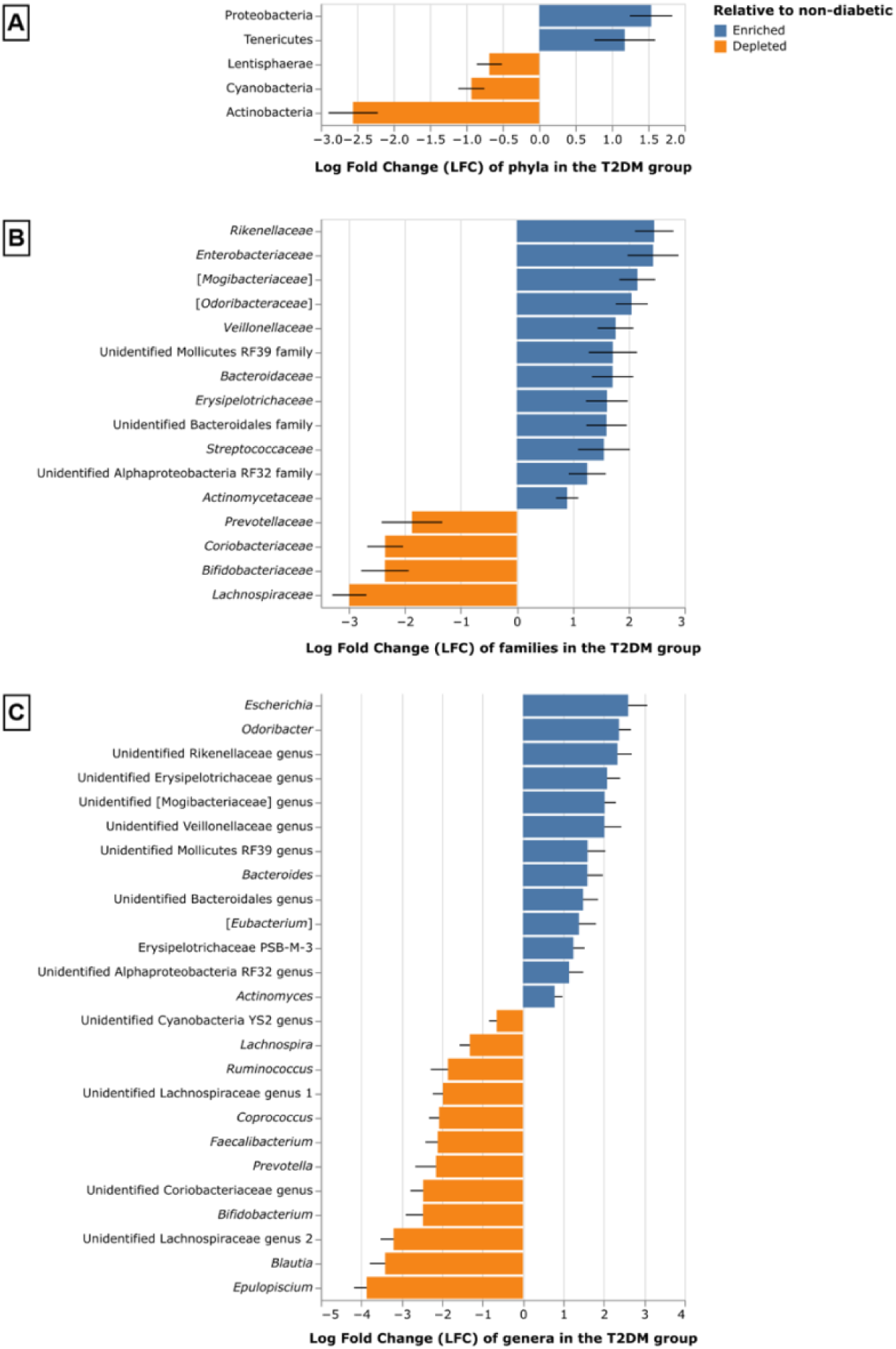
Differentially abundant taxa in T2DM patients compared to non-diabetic individuals at phylum (A), family (B) and genus (C) level. Only taxa with P-values < 0.05 after multiple test correction have been shown. Taxa in square brackets are awaiting reclassification and taxa listed as “Unidentified” were classified to the specified level but had no annotation available in the database.

### 3.6. Alpha diversity metrics in T2DM patients and non-diabetic individuals

The comparison of alpha diversity metrics between the diabetic and non-diabetic groups revealed noteworthy findings. Statistically significant differences were observed between individuals with T2DM and those without diabetes across all alpha diversity metrics, including the Shannon index, Faith’s PD, Pielou’s evenness index and observed features. All four measures indicated significant differences (P < 0.001), with non-diabetic individuals displaying higher median values in comparison to T2DM patients (Table 3). Figure 6 shows the distribution of alpha diversity metrics in T2DM and non-diabetic groups.

**Figure 6.**
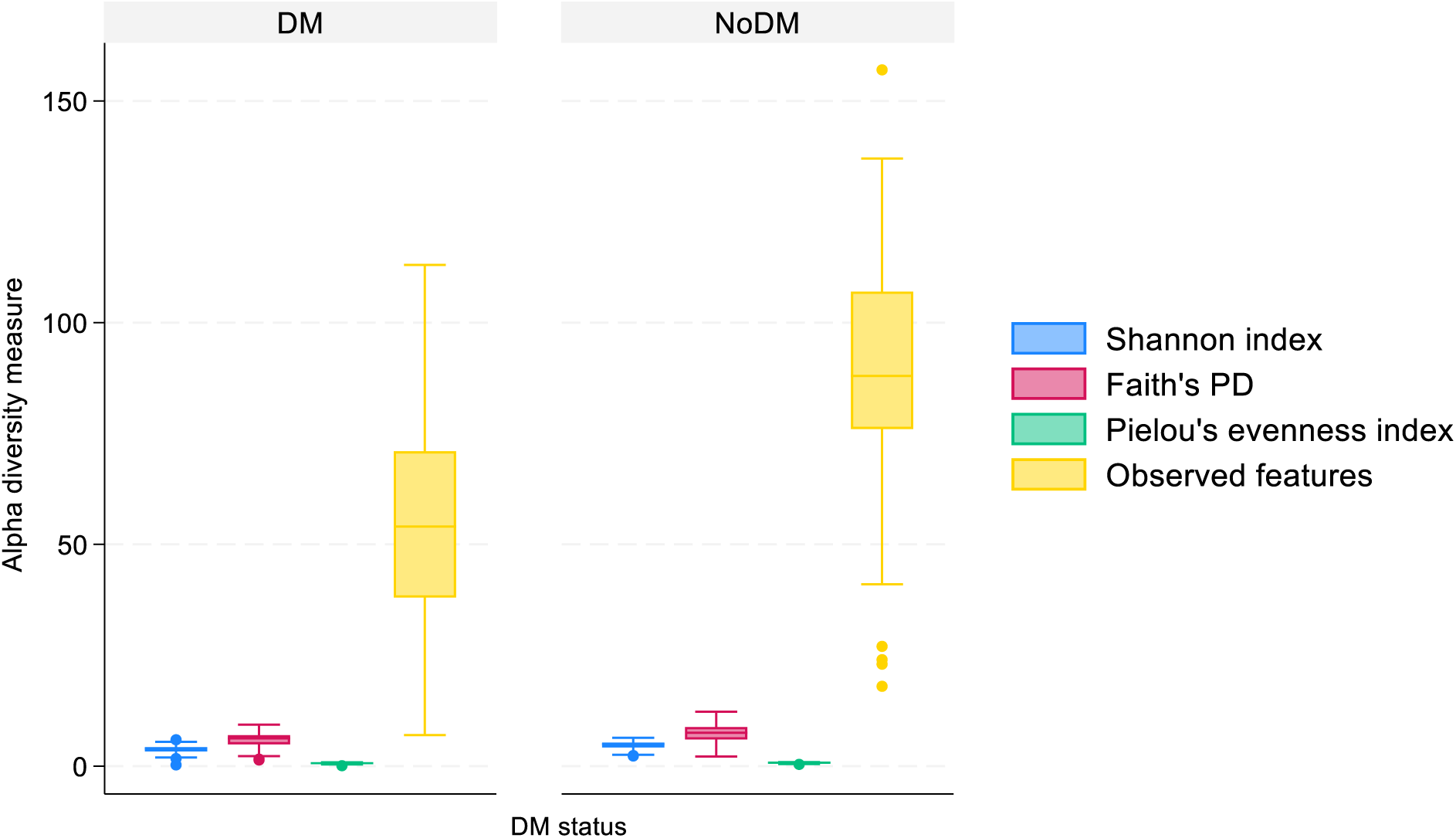
Distribution of alpha diversity metrics in T2DM and non-diabetic groups. DM: diabetes mellitus: No DM: no diabetes mellitus.

**Table 3.** Comparison of alpha diversity metrics between T2DM patients and non-diabetic individuals.

| Alpha diversity metrics | DM group | No DM group | P |
| --- | --- | --- | --- |
| Shannon index | 3.95 (0.98) | 4.98 (1.11) | <0.001 |
| Faith’s PD | 6.26 (2.08) | 7.52 (2.82) | <0.001 |
| Pielou’s evenness index | 0.67 (0.15) | 0.77 (0.14) | <0.001 |
| Observed features | 54 (33) | 88 (31) | <0.001 |

Alpha diversity metrics presented as median and interquartile range. DM: diabetes mellitus; No DM: no diabetes mellitus.

### 3.7. Beta diversity metrics in T2DM patients and non-diabetic individuals

The permutational multivariate analysis of variance using the Bray-Curtis distance, unweighted UniFrac distance and weighted UniFrac distance matrices revealed that samples from the T2DM group were significantly different from the ones from the non-diabetic group (P = 0.001 for all matrices) (Figure 7). The Bray-Curtis clustering pattern accounted for 15.66% and 10.74% of the total variance along the axis 1 and axis 2, respectively. Similarly, the unweighted UniFrac pattern explained 14.14% and 10.45% of the total variance along axis 1 and axis 2. The weighted UniFrac clustering pattern reflected 27.73% and 17.24% of the total variance along axis 1 and axis 2 (Figure 7).

**Figure 7.**
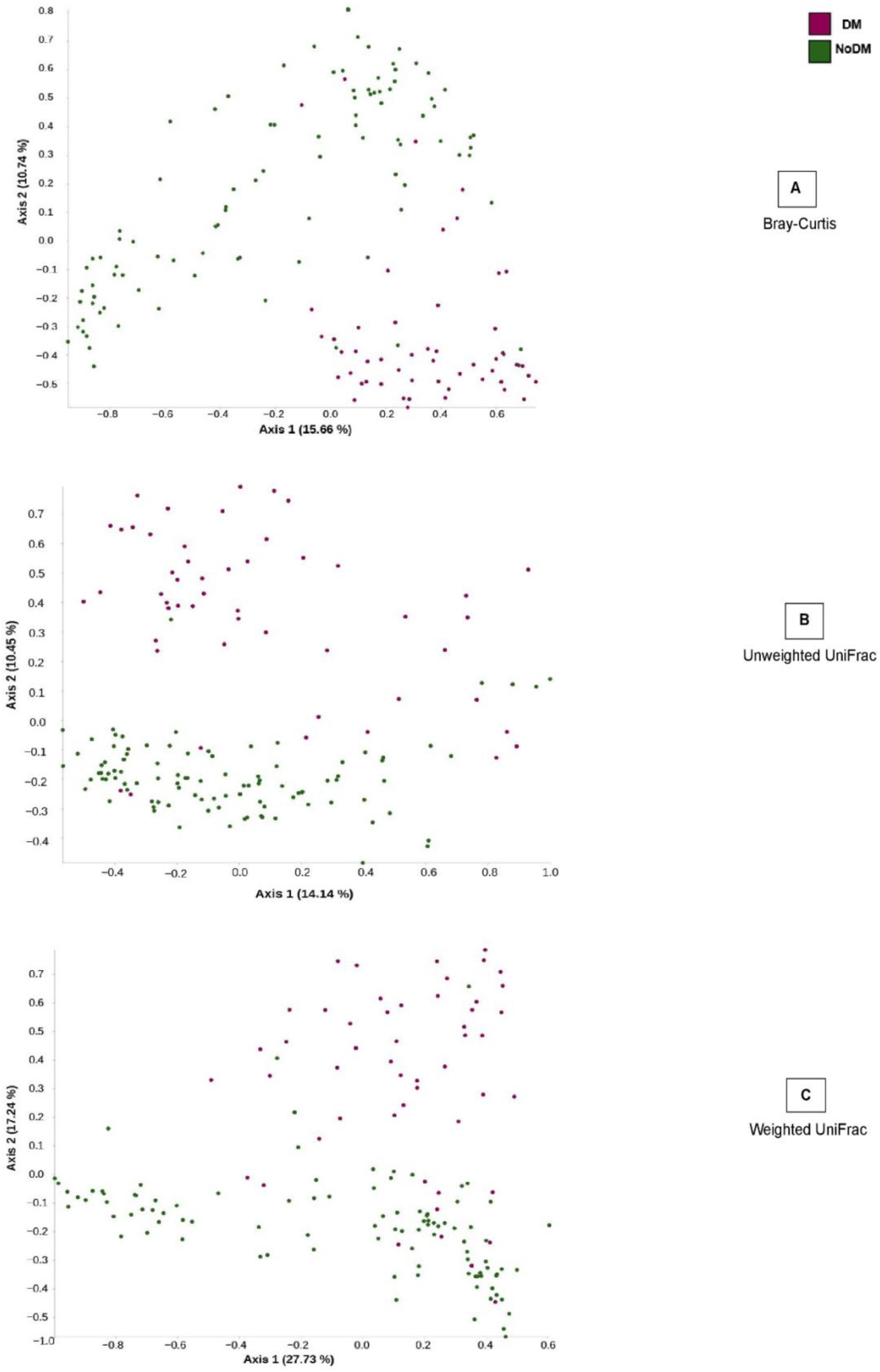
Principal coordinate analysis (PCoA) diagram based on Bray-Curtis (A), unweighted UniFrac (B), and weighted UniFrac (C) distances. Each point on the plots corresponds to a single sample within its respective group. DM: diabetes mellitus: No DM: no diabetes mellitus.

### 3.8. Firmicutes/Bacteroidota (F/B) ratio

The range extended from a minimum ratio of 0.04 to a maximum of 13.4. Specifically, within the T2DM group, the median ratio was 1.4, with an IQR of 1.2. On the other hand, the non-diabetic group displayed a median ratio of 1.5, with an IQR of 2.4 (Table S2). The F/B ratios did not exhibit a statistically significant difference between the T2DM and non-diabetic groups, P = 0.7753.

### 3.9. Relationship of gut microbiota profiles with inflammatory markers, HbA1C, diabetes mellitus status, and type of diabetes mellitus treatment

Binary logistic regression results indicated that Proteobacteria (P = 0.001), *Bacteroidaceae* (P = 0.003), *Erysipelotrichaceae* (P = <0.001), *Enterobacteriaceae* (P = <0.001), *Bacteroides* (P = 0.004), *Holdemanella* (P = 0.001), *Phascolarctobacterium* (P = 0.005), and *Escherichia-Shigella* (P = 0.002) were positively related to the likelihood of falling into the T2DM group. Conversely, the probability of having high levels of *Actinobacteriota* (P = 0.024), *Lachnospiraceae* (P = <0.001), *Ruminococcaceae* (P = 0.003), *Prevotellaceae* (P = 0.001), *Faecalibacterium* (P = <0.001), *Agathobacter* (P = <0.001), *Prevotella_9* (P = 0.002), *Blautia* (P = 0.003), Shannon index (P = <0.001), Faith’s PD (P = <0.001), Pielou’s evenness index (P = <0.001), and observed features (P = <0.001) was more in the non-diabetic group compared to the T2DM group.

Regarding T2DM treatment type, *Lachnospiraceae* exhibited a positive association with the likelihood of belonging to the metformin-using group (P = 0.043). The probability of having high levels of *Phascolarctobacterium* (P = 0.025), Faith’s PD (P = 0.010), and observed features (P = 0.016) was more in the group receiving other T2DM treatments (i.e., insulin, both metformin & insulin, and other unspecified treatments) compared to the metformin-using group.

Correlation analyses indicated a moderate correlation at the phyla level, where Actinobacteriota exhibited a significant negative correlation with HbA1c. Two additional slight associations were noted at the phylum level, consisting of a significant negative correlation between Actinobacteria and CRP, and a significant positive correlation between Proteobacteria and HbA1c (Figure 8).

**Figure 8:**
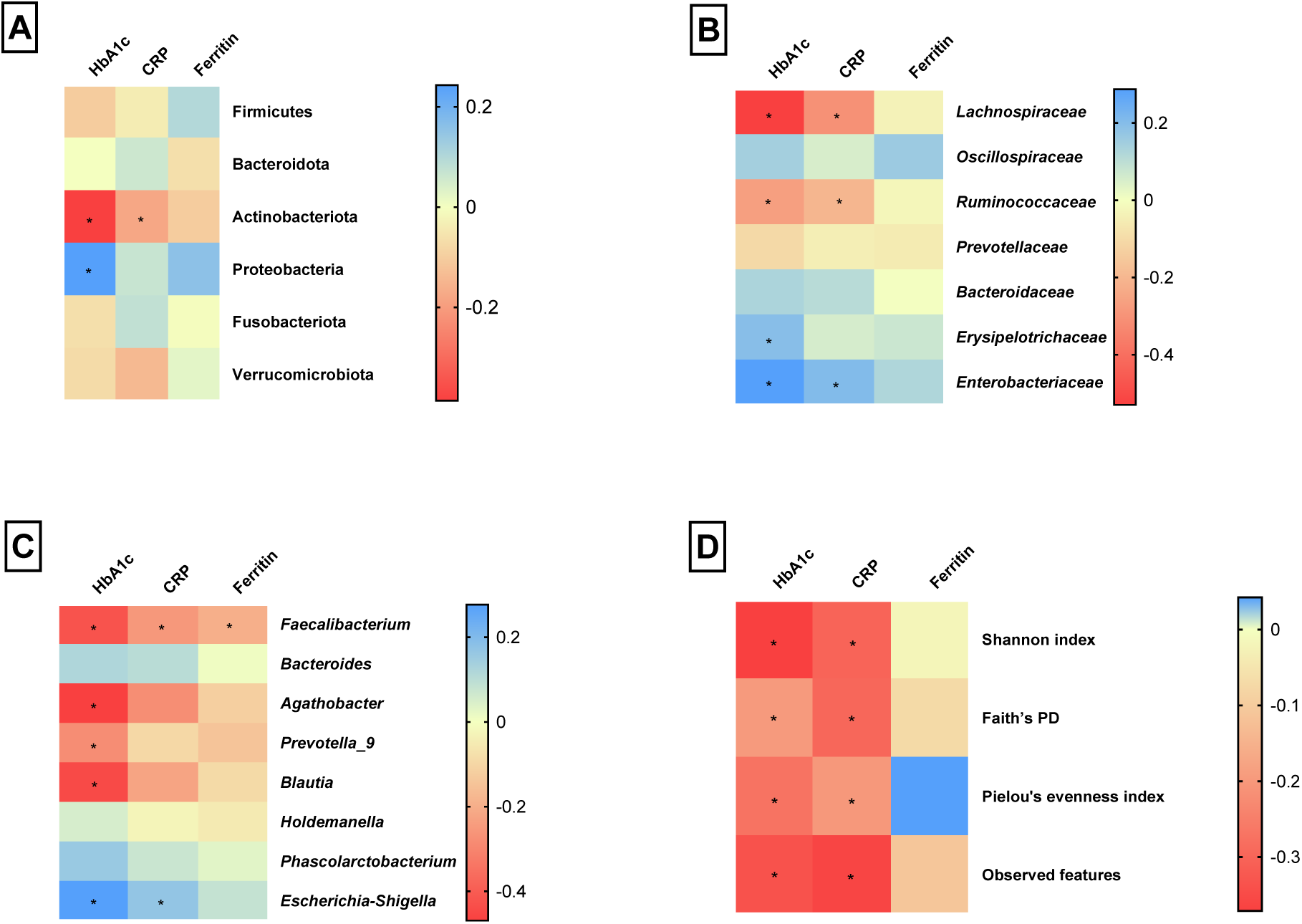
Correlation of HbA1c, CRP and ferritin with relative abundance of the top gut microbes at the phyla, families, and genera and alpha diversity metrics

*Lachnospiraceae* displayed a moderate negative correlation with both HbA1c and CRP. *Ruminococcaceae* showed a moderate negative correlation with HbA1c and a slight negative correlation with CRP. *Erysipelotrichaceae* and *Enterobacteriaceae* exhibited slight positive correlations with HbA1c and CRP, respectively. Additionally, *Enterobacteriaceae* demonstrated a further moderate positive correlation with HbA1c. *Faecalibacterium* displayed a moderate negative association with both HbA1c and CRP, and a slight negative association with ferritin (Figure 8).

*Agathobacter*, *Prevotella_9*, and *Blautia* were moderately negatively correlated with HbA1c. *Escherichia-Shigella* was moderately negatively associated with HbA1c and slightly positively correlated with CRP. All these associations were found to be statistically significant. The Shannon index and Observed features demonstrated moderate negative correlations with both HbA1c and CRP. Faith’s PD and Pielou’s evenness index showed slight negative correlations with HbA1c and CRP, respectively. Additionally, Faith’s PD and Pielou’s evenness index exhibited moderate negative correlations with CRP and HbA1c, respectively (Figure 8).

## 4. Discussion

Investigating the gut microbiota and the features of a balanced microbial community is crucial for understanding the mechanisms through which the gut microbiome influences T2DM. In this study, a comprehensive analysis of gut microbiota composition in T2DM patients and individuals without diabetes was conducted. The findings revealed differences in diversity and the occurrence as well as distribution of phyla, families, and genera between these two groups, offering insights into potential implications for T2DM. Four dominant phyla, Firmicutes, Bacteroidota, Proteobacteria and Actinobacteriota were observed in both groups, which aligns with other gut microbiome studies [28–30]. While previous studies reported comparable abundance of these phyla in both groups [31], our investigation revealed some distinctions.

Proteobacteria demonstrated a higher relative abundance in the T2DM group, emphasizing its potential relevance to diabetes. This supports research conducted by Larsen et al., (2010) [7] and supports the notion of Proteobacteria as a potential microbial indicator of disease [32]. Previous studies have consistently shown an elevated presence of organisms from this phylum in conditions like metabolic disorders and inflammatory bowel disease [32]. The higher relative abundance of Actinobacteria in the non-diabetic group aligns with their recognized role in conducting essential physiological functions within the gastrointestinal tract, such as breaking down resistant starch, generating acetate to support gut barrier function, and contributing to the maturation of the immune system [33].

Additionally, the T2DM group exhibited Fusobacteriota, while the non-diabetic group presented with Verrucomicrobiota. Fusobacteriota encompasses a group of bacteria known for their association with various human diseases, including colorectal cancer [34]. Therefore, its prevalence in the T2DM group may suggest a role in the pathogenesis or progression of diabetes. Verrucomicrobiota is a phylum that includes bacteria known for their mucin-degrading capabilities [35] and their potential to influence gut barrier function and metabolic health. Research has indicated that certain species within Verrucomicrobiota, such as *Akkermansia muciniphila*, may have beneficial effects on metabolic disorders, including improving glucose metabolism and reducing inflammation [36].

Examining the family-level composition, this study shows that in both the T2DM and non-diabetic groups, the predominant families consist of *Bacteroidaceae*, *Prevotellaceae*, and *Lachnospiraceae*. Notably, *Bacteroidaceae* exhibited greater relative and differential abundance in diabetic patients. *Bacteroidetes* species within the *Bacteroidaceae* family have been observed to generate beneficial bacterial gut metabolites such as succinic acid, acetic acid, and, in certain instances, propionic acid as their primary end-products [37]. *Prevotellaceae* and *Lachnospiraceae* were significantly enriched in non-diabetic individuals. The increased occurrence of *Prevotellaceae* in non-diabetic individuals is expected as they have been associated with promoting anti-inflammatory effects and maintaining the integrity of the intestinal barrier [38].

The dominance of *Lachnospiraceae* in the non-diabetic group is in keeping with previously documented research [39]. Moreover, *Lachnospiraceae* was reported by Doumatey et al., (2020) [10] as the most prevalent family in healthy controls. However, Wang *et al.*, (2017) [40] documented the dominance of *Lachnospiraceae* in both diabetic and non-diabetic groups. *Lachnospiraceae* species have been documented to break down starch and other sugars through hydrolysis, resulting in the production of butyrate and other SCFAs [41]. These SCFAs are responsible for beneficial roles that result in reduced levels of inflammatory markers [41].

*Enterobacteriaceae* and *Erysipelotrichaceae* were also among the top families observed in T2DM, and were also shown to be significantly enriched by differential abundance analysis. The increased occurrence of *Enterobacteriaceae* in T2DM has been previously documented [42] and was shown to be associated with imbalances in the gut microbiota and inflammation [43]. Limited information exists regarding the associations of the *Erysipelotrichaceae* family with T2DM. However, available data implies a potential link between *Erysipelotrichaceae* and metabolic disorders in humans [44]. Moreover, members of the *Erysipelotrichaceae* bacterial family seem to provoke a strong immune response and have the potential to thrive even following treatment with broad-spectrum antibiotics [45]. The enrichment of opportunistic pathogens, such as *Escherichia* and *Odoribacter*, in the T2DM group is also concerning. A depletion of genera associated with good health in the diabetic patients, such as *Faecalibacterium* [46] and *Bifidobacterium* [47], was noted. All these findings can negatively affect the management of T2DM, contributing an additional burden to their existing health challenges.

Among individuals without diabetes, *Ruminococcaceae* and *Oscillospiraceae* were also identified as prominent families. The *Oscillospiraceae* family demonstrates an inverse correlation with body mass index. Furthermore, a reduction in *Oscillospiraceae* family members was observed in obese children [48]. In addition, certain members of the *Oscillospiraceae* family generate acetate [49]. Additionally, *Ruminococcaceae* is crucial for preserving gut health as it has the capability to produce butyrate and other SCFAs [50]. A reduced presence of *Ruminococcaceae* has been associated with various inflammatory bowel diseases [50].

The genus level was also studied and the most abundant genus in T2DM patients was *Bacteroides*. The results of the present study agree with those of Wu et al., (2010) [51]. However, Zhao et al., (2020) [52] and Polidori et al., (2022) [53] demonstrated a decreased abundance of *Bacteroides* in their studies. The contradictory results ascertain the complex nature of the gut microbiota and display how different populations can acquire different gut microbiota, due to various factors such as geographical location, diet, exercise [54] and other lifestyle practices. Furthermore, similar to the T2DM group, *Bacteroides* were abundant in the individuals without diabetes. The contribution of various *Bacteroides* species to human health and diseases has been shown to be controversial [55]. *Bacteroides* contribute to the development of intestinal dysfunctions, yet they have demonstrated a level of beneficial influence to the extent that they are considered for use as next-generation probiotics, owing to their potential in enhancing host health [55].

In addition, *Escherichia-Shigella* and *Holdemanella* were also abundant in the T2DM group. The increased abundance of *Escherichia-Shigella* in T2DM was displayed in literature as well [52,56]. Conversely, Huang et al., (2021) [29] reported that a lower abundance of *Escherichia-Shigella* was observed in patients with diabetes. *Escherichia-Shigella* is recognized as a pro-inflammatory microorganism and is associated with gut dysbiosis, potentially contributing to intestinal damage, and influencing amino acid metabolism [57]. Unlike *Escherichia-Shigella*, the *Holdemanella* genus has been linked to both beneficial and detrimental effects on human health. *Holdemanella* was shown to play a crucial role in safeguarding against numerous diseases and various pathogens [58]. However, contradictory findings were documented in previous research, indicating an enrichment of *Holdemanella* in various health conditions such as colorectal cancer, chronic kidney disease, and autism spectrum disorder [59].

The *Phascolarctobacterium* and *Blautia* were also identified as some of the main genera contributing to the overall microbial diversity of T2DM patients in the present study. A recent study by Li et al., (2022) [60] reported a decreased abundance of the *Phascolarctobacterium* family, which is contradictory to the results of the present study. Moreover, in individuals with obesity and elevated BMI, Naderpoor *et al*., (2019) [61] observed a notable positive correlation between *Phascolarctobacterium* and insulin sensitivity. It is crucial to highlight this point since insulin resistance serves as a key connection between obesity and T2DM [61]. Another family that was commonly found in both individuals with T2DM and those without diabetes was *Blautia*. Literature has demonstrated *Blautia* to exhibit a positive correlation with T2DM [6]. This contrasts with the findings of the differential abundance analysis performed in this study, which revealed a depletion in *Blautia* abundance in patients with T2DM compared to individuals without diabetes. *Blautia* has received significant attention due to its role in mitigating inflammatory and metabolic diseases, as well as its antibacterial effects against specific microorganisms [62]. Recent studies highlighted the correlation between the composition and fluctuations in the *Blautia* population in the gut with various factors, including but not limited to geographical location, diet, health, disease status, and other physiological conditions [62]. Additionally, this genus has been identified to play a specific role in biotransformation and interactions with other intestinal microorganisms [62].

Looking at non-diabetic individuals, *Prevotella_9* was the most prevalent, representing most of the non-diabetic group microbial community. *Prevotella_9* is linked to a plant-based, low-fat diet and serves as important bacterial constituents during the maturation of the human gut microbiota from infancy to young adulthood [58]. According to Tao et al., (2019) [63], their similar findings displayed a significant reduction in the presence of *Prevotella_9* within the group with T2DM. *Prevotella_9* exerts positive effects on intestinal disorders through the regulation of the immune system and maintenance of intestinal homeostasis [64]. Moreover, the *Prevotella_9* genus is recognized for its role in generating propionate [65]. Propionate, a significant by-product of microbial fermentation in the human gut, is believed to have health effects [66] and seems to play a crucial role in the positive effects of the gut microbiome [67]. *Agathobacter* and *Faecalibacterium* were also among the leading genera observed in individuals without diabetes, although only the latter was significantly enriched in this group, when considering differential abundance. Siptroth et al., (2023) [39] identified a decline in the *Agathobacter* genus within the T2DM group in their study, which aligns with our study. *Agathobacter* is known to generate SCFAs, including butyrate, and is associated with beneficial intestinal flora [39]. A decrease in this genus leads to reduced proportion of SCFAs in the body, therefore promoting conditions like obesity [39]. Consequently, this genus could serve as a component in a profile for detecting T2DM [39]. However, further experimental studies are necessary to understand the relationship between the *Agathobacter* genus and the development of T2DM [39].

In about half of the microbiome studies related to T2DM reviewed by Gurung et al., (2020) [6], a reduction in *Faecalibacterium* levels was identified in T2DM patients, indicating a possible significance beyond its biomarker role. Other research investigations have emphasised the association between inflammatory conditions and diminished levels of *Faecalibacterium* [46]. Moreover, the abundance of this genus is regarded as indicative, to some extent, of the state of intestinal health since patients with gastrointestinal disorders often display lower levels of *Faecalibacterium* [46].

Additional evidence supporting the distinction between the gut microbiota of patients with T2DM and individuals without diabetes is provided by the alpha and beta diversity findings. The current study indicates that there are significant differences in the microbial diversity between patients with T2DM and individuals without diabetes. The non-diabetic group tends to have a higher diversity, both in terms of the number of observed features and the evenness of the microbial community, as indicated by the alpha diversity metrics. Some studies also reported that diversity and richness were increased in non-diabetic individuals as compared to diabetic patients [63,68]. Contrasting the present study, Doumatey et al., (2020) [10] found that the alpha-diversity indices were significantly greater in T2DM cases compared to non-diabetic controls.

This study further employed three different distance metrics (Bray-Curtis, un-weighted UniFrac, and weighted UniFrac) to assess dissimilarities in the microbial communities. All the beta diversity metrics showed significant differences between the T2DM and non-diabetic group. Que et al., (2021) [69] reported similar outcomes. Conversely, Almugadam *et al.*, (2020) [68] reported contrasting findings.

Following the assessment of the F/B ratio, our findings indicate no significant difference in F/B ratios between the T2DM and non-diabetic groups. However, Salamon *et al*., (2018) [70] observed conflicting findings. Moreover, Kusnadi et al., (2023) [71] conducted a systematic review, with studies comparing the F/B ratio in patients with T2DM and individuals with a normal glycaemic index. Among these studies, half of them indicated that the F/B ratio was higher in the T2DM group compared to the control group. While the alterations in F/B ratio patterns may not consistently follow a specific trend in patients with diabetes, an altered F/B ratio indicates a dysbiotic gut microbiota [72].

The identified positive associations of specific bacteria in individuals with T2DM may imply a potential involvement of these microbes in the onset or progression of T2DM. Conversely, the negative associations with certain bacteria in the T2DM group could indicate a potential protective role or a more favorable composition of the gut microbiome in non-diabetic individuals. Furthermore, the observed association between *Lachnospiraceae* and metformin usage suggests a potential impact of this medication on the composition of gut microbiota. However, a larger sample size is required to confirm this finding.

The present study also revealed that most patients with T2DM (44%) demonstrated a moderate increase in CRP levels, and among female T2DM patients, 20% exhibited elevated ferritin levels. In comparison, only 2% of the non-diabetic female group had high blood ferritin levels, and the majority (55%) of the non-diabetic group displayed normal CRP levels.

Furthermore, there were correlations of CRP with bacterial taxa such *Lachnospiraceae* and *Faecalibacterium*, as well as all alpha diversity metrics studied in the present study. Conversely, a study examining the correlation between gut microbiota and inflammatory markers in patients with both obesity and T2DM was conducted and found no observed association [73]. Only a limited number of studies have examined a wide range of inflammatory markers in relation to the gut microbiota [20]. Therefore, there is a need for additional studies of a similar nature.

## 5. Limitations of the study

In this study, no negative controls were sequenced. Nevertheless, a negative control was used during the PCR stage to evaluate the presence of contaminations. There were fewer than expected species level classifications in this study, which was most likely a result of the information available in the databases used within DADA2 and QIIME2 with PacBio CCS. One of the other primary limitations of this study is the small sample size (150), which may restrict the generalisability of our findings to the broader South African population. The study’s focus on a specific geographical region might limit the applicability of our results to populations with distinct genetic, cultural, and environmental backgrounds. Variations in individual dietary habits [74] and physical activity [75], which were not comprehensively assessed, may contribute to the observed differences in gut microbiota and could confound the association with T2DM. Moreover, insufficient details about concurrent health conditions and lifestyle decisions, like tobacco, alcohol, and drug use, could potentially influence the study outcomes. Body mass index is a key anthropometric measure commonly associated with T2DM and its omission from the current study may introduce potential confounding, as variations in BMI among individuals, irrespective of diabetes status, could contribute to differences in the gut microbial landscape. Information about T2DM duration was not available, consequently, this was not uniformly controlled. This variation may introduce additional complexities when attributing changes in gut microbiota composition to diabetes status. Other drugs in addition to antidiabetic medications such as metformin may exert direct or indirect effects on microbial communities, and their impact on our findings remains a noteworthy limitation. Moreover, A weakness of this study is that the sample size of 51 T2DM patients is too small to support detailed sub-analyses, such as comparing those on metformin with those not on the medication. While the study focused on taxonomic composition, it did not delve deeply into the functional aspects of the gut microbiota. Investigating the metabolic functions and interactions among microbial species could provide a more nuanced understanding of the microbial contributions to T2DM.

## 6. Conclusions

The examination of gut microbiota in patients with T2DM and individuals without diabetes revealed clear differences. Both groups highlighted microbial phyla, families, and genera associated with both health and disease. However, a notable observation was the higher abundance of harmful taxa among T2DM patients compared to non-diabetic individuals, who exhibited a greater concentration of beneficial taxa. Additionally, the comparison of alpha and beta diversity metrics between the two groups yielded statistically significant differences. Furthermore, our findings indicated some evident associations between inflammatory markers and diversity/richness. The findings contribute to the growing understanding of the complex relationship between the gut microbiota and T2DM. Insights gained from studying gut microbiota in the South African T2DM population have the potential to inform the public and policy makers about simple and affordable supplementary therapeutic measures that can be considered for management of T2DM. Although the present study is observational in nature, it is perceived as an important foundation to encourage the conduct of experimental studies that can employ affordable probiotic-rich diets that could potentially improve the gut microbiota.

## Ethics approval and consent to participate

Ethics approval was attained from the Stellenbosch University health research ethics committee (HREC), HREC Reference No: S22/05/009_COVID-19 (PhD). The research adhered to the principles outlined in the Declaration of Helsinki and guidelines established by the HREC. Participants were recruited from the Dr. George Mukhari Hospital (DGMAH) and nearby townships and provided their consent to participate in the study.

## Consent for publication

Not applicable

## Availability of data and materials

16S rRNA sequencing data can be found here: https://dataview.ncbi.nlm.nih.gov/object/PRJNA1064418. Submission ID: SUB14150586, BioProject ID: PRJNA1064418.

## Author’s contributions

SMP came up with the study concept and wrote the first draft of the manuscript. PSN, BC-M, and SM helped to develop the study and revise the manuscript. JTNN, AS and KN performed bioinformatics analysis and revised the manuscript. All authors approved the last version of the manuscript.

## Acknowledgements

We express our gratitude to Sister Thenjiwe and the Phlebotomists (Portia, Tebogo, Lindo, Rose, and Mmasina), Doctors Matladi, Maluleke, Mohale, Malapermala, and Mahlangu, as well as the student Scientists (Scelo, Nelly, Luthando, and Bridgette) for their invaluable assistance in sample and data collection. Special thanks to Mr. Hleki Rikhotso for his assistance in navigating the homes of the patients and collecting samples from house to house, alongside the principal investigator (SMP). We would like to extend our heartfelt appreciation to all the study participants for their time, patience, and cooperation.

---

Alpha diversity plots

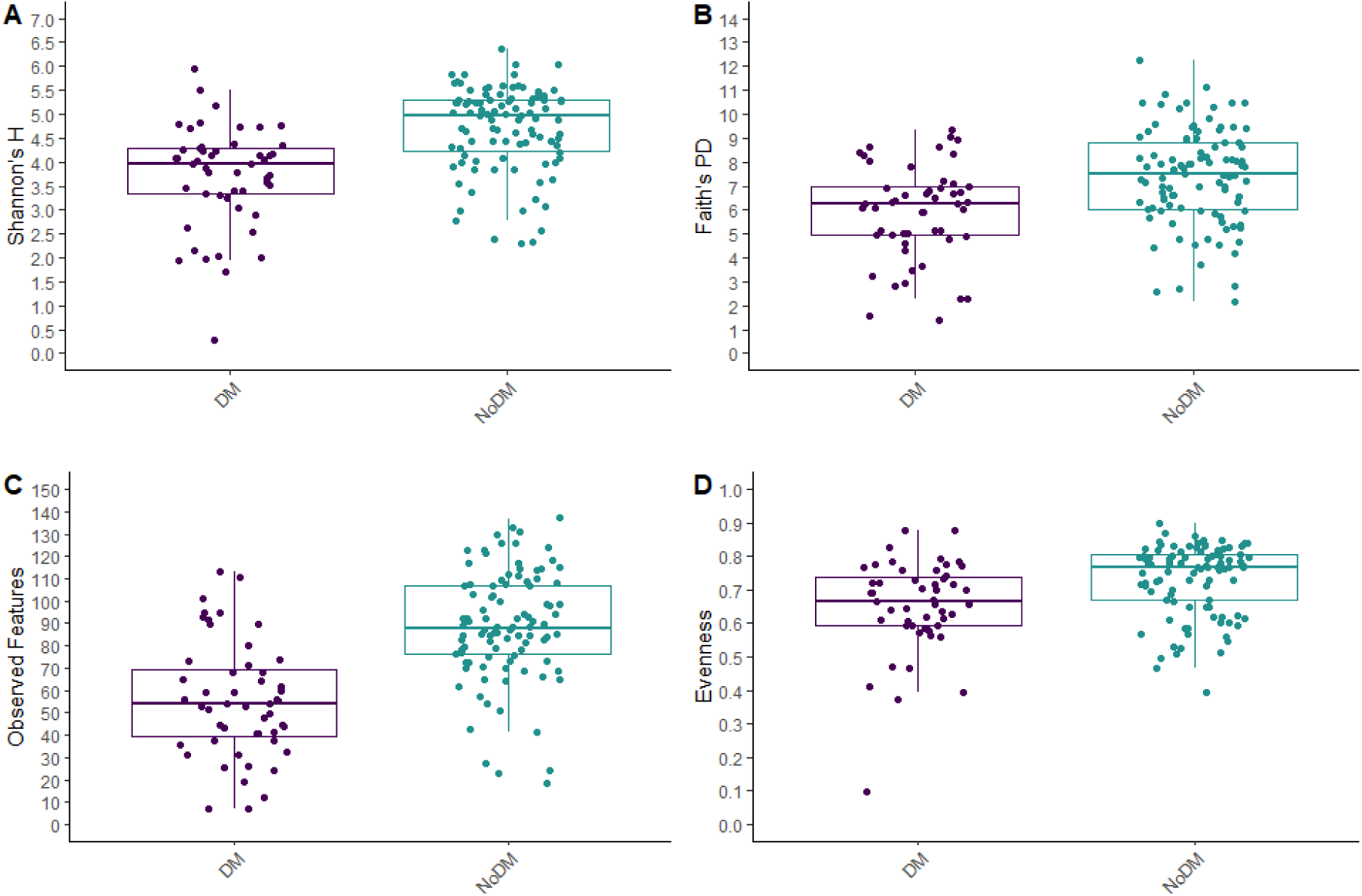

Bray Curtis with centroids (colour blind friendly - R values need to be added)

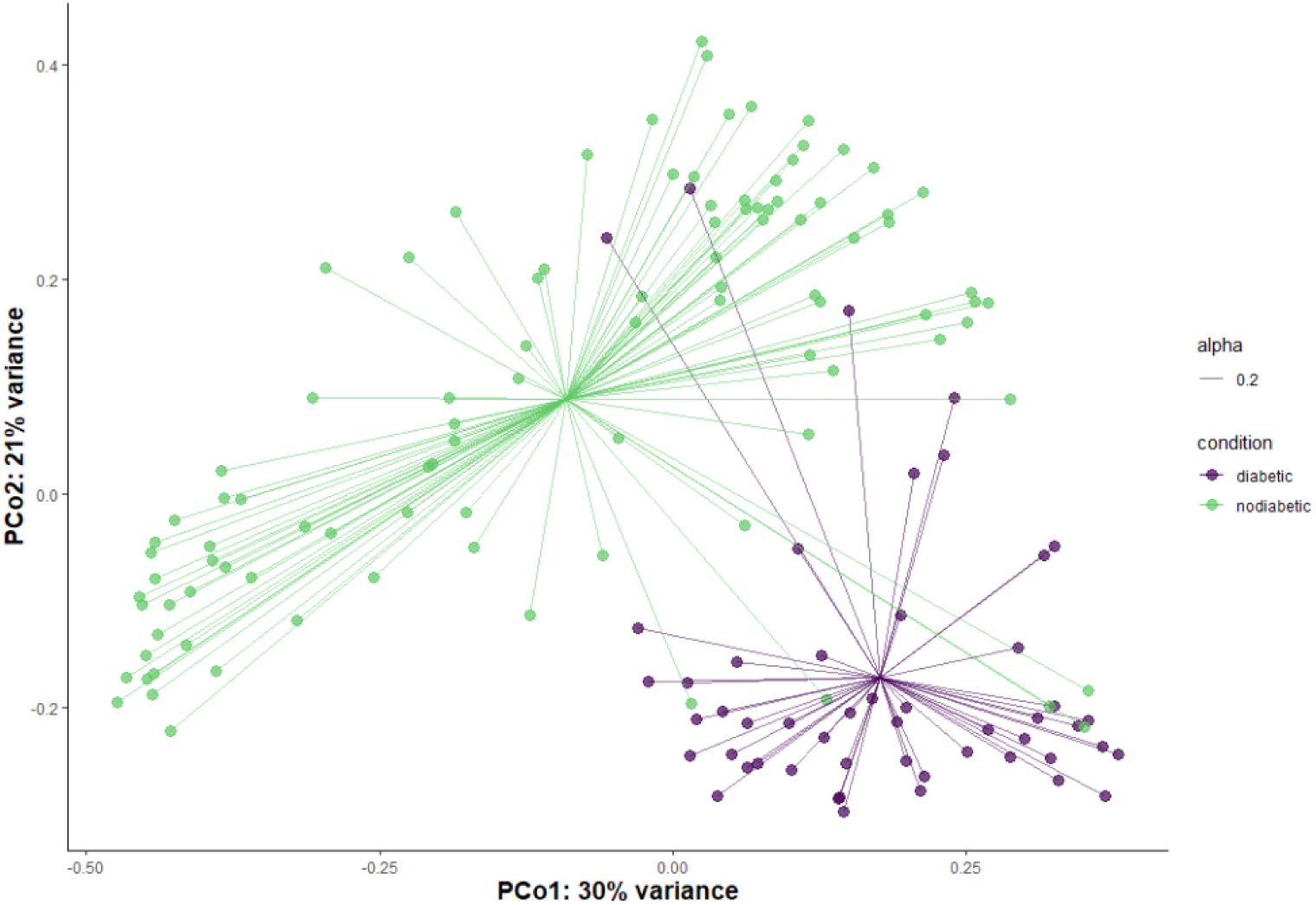

Bray Curtis with ellipses

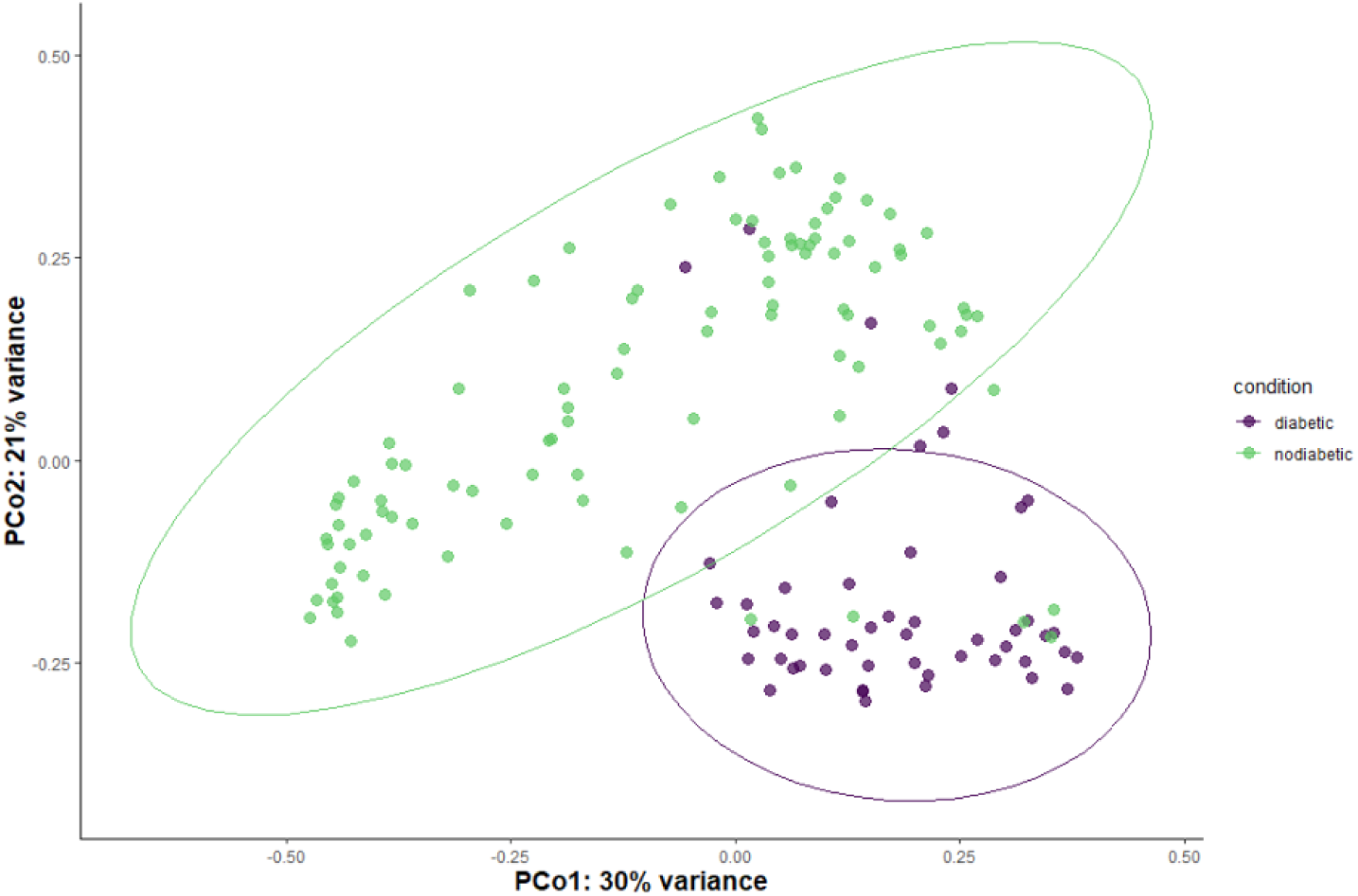

## References

1. Thursby E, Juge N. Introduction to the human gut microbiota. Biochem J. 2017;474(11):1823–36.

2. Valdes AM, Walter J, Segal E, Spector TD. Role of the gut microbiota in nutrition and health. BMJ (Online). 2018; 361:k2179:36–44.

3. Peter PI, Steinberg WJ, van Rooyen C, Botes J. Type 2 diabetes mellitus patients’ knowledge, attitude and practice of lifestyle modifications. Health SA. 2022; 27:1921.

4. Ong KL, Stafford LK, McLaughlin SA, Boyko EJ, Vollset SE, Smith AE, et al. Global, regional, and national burden of diabetes from 1990 to 2021, with projections of prevalence to 2050: a systematic analysis for the Global Burden of Disease Study 2021. Lancet. 2023;402(10397):203–34.

5. Chetty L, Govender N, Govender GM, Reddy P. Demographic stratification of type 2 diabetes and comorbidities in district healthcare in Kwazulu-Natal. S Afr Fam Pract (2004). 2021; 63(1):e1–e9.

6. Gurung M, Li Z, You H, Rodrigues R, Jump DB, Morgun A, et al. Role of gut microbiota in type 2 diabetes pathophysiology. EBioMedicine. 2020;51:1–9.

7. Larsen N, Vogensen FK, Van Den Berg FWJ, Nielsen DS, Andreasen AS, Pedersen BK, et al. Gut microbiota in human adults with type 2 diabetes differs from non-diabetic adults. PLoS ONE. 2010; 5(2):e9085.

8. Sedighi M, Razavi S, Navab-Moghadam F, Khamseh ME, Alaei-Shahmiri F, Mehrtash A, et al. Comparison of gut microbiota in adult patients with type 2 diabetes and healthy individuals. Microb Pathog. 2017;111:362–9.

9. Zhao L, Lou H, Peng Y, Chen S, Zhang Y, Li X. Comprehensive relationships between gut microbiome and faecal metabolome in individuals with type 2 diabetes and its complications. Endocrine. 2019;66(3):526–37.

10. Doumatey AP, Adeyemo A, Zhou J, Lei L, Adebamowo SN, Adebamowo C, et al. Gut Microbiome Profiles Are Associated With Type 2 Diabetes in Urban Africans. Front Cell Infect Microbiol. 2020; 10:63.

11. Allali I, Abotsi RE, Tow LAh, Thabane L, Zar HJ, Mulder NM, et al. Human microbiota research in Africa: a systematic review reveals gaps and priorities for future research. Microbiome. 2021;9(294).

12. Lu J, Ma KL, Ruan XZ. Dysbiosis of Gut Microbiota Contributes to the Development of Diabetes Mellitus. Infec Microb Dis. 2019; 1(2): 43–48.

13. Violi F, Nocella C, Bartimoccia S, Castellani V, Carnevale R, Pignatelli P, et al. Gut dysbiosis-derived low-grade endotoxemia: A common basis for liver and cardiovascular disease. Kardiol Pol. 2023;81(6):563–571.

14. Ciesielska A, Matyjek M, Kwiatkowska K. TLR4 and CD14 trafficking and its influence on LPS-induced pro-inflammatory signaling. Cell Mol Life Sci. 2021;78(4):1233–1261.

15. Bashir H, Majid S, Khan MS, Bhat MH, Hamid R, Ashraf R, et al. Inter-relationship of Pro- and Anti-inflammatory Biomarkers with the development of Type 2 Diabetes Mellitus. Heliyon. 2022; 8(11):e11329.

16. Rehman K, Akash MSH. Mechanisms of inflammatory responses and development of insulin resistance: How are they interlinked? J Biomed Sci. 2016;23(1):1–18.

17. Kim MH, Kim HN, Choi WS. The association between subclinical inflammation and abnormal glucose and lipid metabolisms in normal-weight Korean individuals. Nutr Metab Cardiovasc Dis. 2018;28(11):1106–13.

18. Dludla P V, Mabhida SE, Ziqubu K, Nkambule BB, Mazibuko-Mbeje SE, Hanser S, et al. Pancreatic β-cell dysfunction in type 2 diabetes: Implications of inflammation and oxidative stress. World J Diabetes. 2023;14(3):130–46.

19. Deleu S, Machiels K, Raes J, Verbeke K, Vermeire S. Short chain fatty acids and its producing organisms: An overlooked therapy for IBD? EBioMedicine. 2021;66:10329.

20. Bander Z Al, Nitert MD, Mousa A, Naderpoor N. The gut microbiota and inflammation: An overview. Int J Environ Res Public Health. 2020;17(20):1–22.

21. Lau WL, Tran T, Rhee CM, Kalantar-Zadeh K, Vaziri ND. Diabetes and the Gut Microbiome. Semin Nephrol. 2021; 41(2):104–113.

22. Di Tommaso P, Chatzou M, Floden EW, Barja PP, Palumbo E, Notredame C. Nextflow enables reproducible computational workflows. Nat Biotechnol. 2017;35(4):316–9.

23. Ewels PA, Peltzer A, Fillinger S, Patel H, Alneberg J, Wilm A, et al. The nf-core framework for community-curated bioinformatics pipelines. Nat Biotechnol. 2020;38(3):276–8.

24. Straub D, Blackwell N, Langarica-Fuentes A, Peltzer A, Nahnsen S, Kleindienst S. Interpretations of Environmental Microbial Community Studies Are Biased by the Selected 16S rRNA (Gene) Amplicon Sequencing Pipeline. Front Microbiol. 2020;11:550420.

25. Ewels P, Magnusson M, Lundin S, Käller M. MultiQC: Summarize analysis results for multiple tools and samples in a single report. Bioinformatics. 2016;32(19):3047–8.

26. Sherwani SI, Khan HA, Ekhzaimy A, Masood A, Sakharkar MK. Significance of HbA1c test in diagnosis and prognosis of diabetic patients. Biomark Insights. 2016;11:95–104.

27. Nehring SM, Goyal A, Patel BC. C Reactive Protein. (Updated 2023 Jul 10). In: StatPearls (Internet). Treasure Island (FL): StatPearls Publishing; 2024 Jan-. Available from: https://www.ncbi.nlm.nih.gov/books/NBK441843/

28. Cunningham AL, Stephens JW, Harris DA. Gut microbiota influence in type 2 diabetes mellitus (T2DM). Gut Pathog. 2021;13(50).

29. Huang Y, Wang Z, Ma H, Ji S, Chen Z, Cui Z, et al. Dysbiosis and Implication of the Gut Microbiota in Diabetic Retinopathy. Front Cell Infect Microbiol. 2021;11:646348.

30. Wongsurawat T, Sutheeworapong S, Jenjaroenpun P, Charoensiddhi S, Khoiri AN, Topanurak S, et al. Microbiome analysis of thai traditional fermented soybeans reveals short-chain fatty acid-associated bacterial taxa. Sci Rep. 2023; 13(1):7573.

31. Das T, Jayasudha R, Chakravarthy SK, Prashanthi GS, Bhargava A, Tyagi M, et al. Alterations in the gut bacterial microbiome in people with type 2 diabetes mellitus and diabetic retinopathy. Sci Rep. 2021;11(1):2738.

32. Rizzatti G, Lopetuso LR, Gibiino G, Binda C, Gasbarrini A. Proteobacteria: A common factor in human diseases. Biomed Res Int. 2017;2017:9351507.

33. Binda C, Lopetuso LR, Rizzatti G, Gibiino G, Cennamo V, Gasbarrini A. Actinobacteria: A relevant minority for the maintenance of gut homeostasis. Dig Liver Dis; 2018. p. 421–8.

34. Kelly D, Yang L, Pei Z. Gut Microbiota, Fusobacteria, and Colorectal Cancer. Diseases. 2018;6(4):109.

35. Bielka W, Przezak A, Pawlik A. The role of the gut microbiota in the pathogenesis of diabetes. IntJ Mol Sci. 2022;23(1):480.

36. Li J, Yang G, Zhang Q, Liu Z, Jiang X, Xin Y. Function of Akkermansia muciniphila in type 2 diabetes and related diseases. Front Microbiol. 2023;14:1172400.

37. Rajilić-Stojanović M, de Vos WM. The first 1000 cultured species of the human gastrointestinal microbiota. EMS Microbiol Rev. 2014;38(5):996–1047.

38. Iatcu CO, Steen A, Covasa M. Gut microbiota and complications of type-2 diabetes. Nutrients. 2021; 14(1):166.

39. Siptroth J, Moskalenko O, Krumbiegel C, Ackermann J, Koch I, Pospisil H. Variation of butyrate production in the gut microbiome in type 2 diabetes patients. Int Microbiol. 2023;26(3):601–610.

40. Wang Y, Luo X, Mao X, Tao Y, Ran X, Zhao H, et al. Gut microbiome analysis of type 2 diabetic patients from the Chinese minority ethnic groups the Uygurs and Kazaks. PLoS One. 2017;12(3):1–15.

41. Vacca M, Celano G, Calabrese FM, Portincasa P, Gobbetti M, De Angelis M. The controversial role of human gut lachnospiraceae. Microorganisms. 2020; 8(4):573.

42. Gradisteanu Pircalabioru G, Chifiriuc MC, Picu A, Petcu LM, Trandafir M, Savu O. Snapshot into the Type-2-Diabetes-Associated Microbiome of a Romanian Cohort. Int J Mol Sci. 2022;23(23):15023.

43. Baldelli V, Scaldaferri F, Putignani L, Del Chierico F. The Role of Enterobacteriaceae in Gut Microbiota Dysbiosis in Inflammatory Bowel Diseases. Microorganisms. 2021;9(4):697.

44. Turpin W, Bedrani L, Espin-Garcia O, Xu W, Silverberg MS, Smith MI, et al. Associations of NOD2 polymorphisms with Erysipelotrichaceae in stool of in healthy first degree relatives of Crohn’s disease subjects. BMC Med Genet. 2020;21(1):204.

45. Kaakoush NO. Insights into the role of Erysipelotrichaceae in the human host. Front Cell Infect Microbiol. 2015;5:84.

46. Martín R, Rios-Covian D, Huillet E, Auger S, Khazaal S, Bermúdez-Humarán LG, et al. Faecalibacterium: a bacterial genus with promising human health applications. FEMS Microbiol Rev. 2023;47(4):fuad039.

47. Hidalgo-Cantabrana C, Delgado S, Ruiz L, Ruas-Madiedo P, Sánchez B, Margolles A. Bifidobacteria and Their Health-Promoting Effects. Microbiol Spectr. 2017 ;5(3).

48. Atzeni A, Bastiaanssen TFS, Cryan JF, Tinahones FJ, Vioque J, Corella D, et al. Taxonomic and Functional Fecal Microbiota Signatures Associated With Insulin Resistance in Non-Diabetic Subjects With Overweight/Obesity Within the Frame of the PREDIMED-Plus Study. Front Endocrinol (Lausanne). 2022;13:804455.

49. Palmnäs-Bedard MSA, Costabile G, Vetrani C, Åberg S, Hjalmarsson Y, Dicksved J, et al. The human gut microbiota and glucose metabolism: a scoping review of key bacteria and the potential role of SCFAs. Am J Clin Nutr. 2022;116(4):862–874.

50. Gu X, Sim JXY, Lee WL, Cui L, Chan YFZ, Chang ED, et al. Gut Ruminococcaceae levels at baseline correlate with risk of antibiotic-associated diarrhea. iScience. 2021;25(1):103644.

51. Wu X, Ma C, Han L, Nawaz M, Gao F, Zhang X, et al. Molecular characterisation of the faecal microbiota in patients with type II diabetes. Curr Microbiol. 2010;61(1):69–78.

52. Zhao X, Zhang Y, Guo R, Yu W, Zhang F, Wu F, et al. The Alteration in Composition and Function of Gut Microbiome in Patients with Type 2 Diabetes. J Diabetes Res. 2020;2020:8842651.

53. Polidori I, Marullo L, Ialongo C, Tomassetti F, Colombo R, di Gaudio F, et al. Characterization of Gut Microbiota Composition in Type 2 Diabetes Patients: A Population-Based Study. Int J Environ Res Public Health. 2022;19(23):15913.

54. Wegierska AE, Charitos IA, Topi S, Potenza MA, Montagnani M, Santacroce L. The Connection Between Physical Exercise and Gut Microbiota: Implications for Competitive Sports Athletes. Sports Med. 2022;52(10):2355–2369.

55. Wang C, Zhao J, Zhang H, Lee YK, Zhai Q, Chen W. Roles of intestinal bacteroides in human health and diseases. Crit Rev Food Sci Nutr. 2021;61(21):3518–3536.

56. Maskarinec G, Raquinio P, Kristal BS, Setiawan VW, Wilkens LR, Franke AA, et al. The gut microbiome and type 2 diabetes status in the Multiethnic Cohort. PLoS One. 2021;16(6):e0250855.

57. Yang L, Xiang Z, Zou J, Zhang Y, Ni Y, Yang J. Comprehensive Analysis of the Relationships Between the Gut Microbiota and Fecal Metabolome in Individuals With Primary Sjogren’s Syndrome by 16S rRNA Sequencing and LC–MS-Based Metabolomics. Front Immunol. 2022;13:874021.

58. De D, Nayak T, Chowdhury S, Dhal PK. Insights of Host Physiological Parameters and Gut Microbiome of Indian Type 2 Diabetic Patients Visualized via Metagenomics and Machine Learning Approaches. Front Microbiol. 2022;13:914124.

59. Zhang C, Liang D, Li X, Liu J, Fan M, Jing M, et al. Characteristics of Gut Microbial Profiles of Offshore Workers and Its Associations With Diet. Front Nutr. 2022;9:904927.

60. Li W, Li L, Yang F, Hu Q, Xiong D. Correlation between gut bacteria Phascolarctobacterium and exogenous metabolite α-linolenic acid in T2DM: a case-control study. Ann Transl Med. 2022;10(19):1056.

61. Naderpoor N, Mousa A, Gomez-Arango LF, Barrett HL, Nitert MD, Courten B De. Faecal microbiota are related to insulin sensitivity and secretion in overweight or obese adults. J Clin Med. 2019;8(4):452.

62. Liu X, Mao B, Gu J, Wu J, Cui S, Wang G, et al. Blautia—a new functional genus with potential probiotic properties? Vol. 13, Gut Microbes. 2021;13(1):1–21.

63. Tao S, Li L, Li L, Liu Y, Ren Q, Shi M, et al. Understanding the gut–kidney axis among biopsy-proven diabetic nephropathy, type 2 diabetes mellitus and healthy controls: an analysis of the gut microbiota composition. Acta Diabetol. 2019;56(5):581–592.

64. Zhao H, Lyu Y, Zhai R, Sun G, Ding X. Metformin Mitigates Sepsis-Related Neuroinflammation via Modulating Gut Microbiota and Metabolites. Front Immunol. 2022;13:797312.

65. Sun QH, Liu ZJ, Zhang L, Wei H, Song LJ, Zhu SW, et al. Sex-based differences in fecal short-chain fatty acid and gut microbiota in irritable bowel syndrome patients. J Dig Dis. 2021;22(5):246–255.

66. Hosseini E, Grootaert C, Verstraete W, Van de Wiele T. Propionate as a health-promoting microbial metabolite in the human gut. Nutr Rev. 2011;69(5):245–58.

67. Blaak EE, Canfora EE, Theis S, Frost G, Groen AK, Mithieux G, et al. Short chain fatty acids in human gut and metabolic health. Benef Microbes. 2020;11(5):411–455.

68. Almugadam BS, Liu Y, Chen SM, Wang CH, Shao CY, Ren BW, et al. Alterations of Gut Microbiota in Type 2 Diabetes Individuals and the Confounding Effect of Antidiabetic Agents. J Diabetes Res. 2020;2020:7253978.

69. Que Y, Cao M, He J, Zhang Q, Chen Q, Yan C, et al. Gut Bacterial Characteristics of Patients With Type 2 Diabetes Mellitus and the Application Potential. Front Immunol. 2021;12:722206.

70. Salamon D, Sroka-Oleksiak A, Kapusta P, Szopa M, Mrozińska S, Ludwig-Słomczyńska AH, et al. Characteristics of the gut microbiota in adult patients with type 1 and 2 diabetes based on the analysis of a fragment of 16S rRNA gene using next-generation sequencing. Pol Arch Intern Med. 2018;128(6):336–343.

71. Kusnadi Y, Saleh MI, Ali Z, Hermansyah H, Murti K, Hafy Z, et al. Firmicutes/Bacteroidetes Ratio of Gut Microbiota and Its Relationships with Clinical Parameters of Type 2 Diabetes Mellitus: A Systematic Review. Open Access Maced J Med Sci (Internet). 2023 Feb. 2 (cited 2024 Apr. 25);11(F):67–72.

72. Liu X, Cheng YW, Shao L, Sun SH, Wu J, Song QH, et al. Gut microbiota dysbiosis in Chinese children with type 1 diabetes mellitus: An observational study. World J Gastroenterol. 2021;27(19):2394–2414.

73. Sugawara Y, Kanazawa A, Aida M, Yoshida Y, Yamashiro Y, Watada H. Association of gut microbiota and inflammatory markers in obese patients with type 2 diabetes mellitus: Post hoc analysis of a synbiotic interventional study. Biosci Microbiota Food Health. 2022;41(3):103–111.

74. Yin P, Zhang C, Du T, Yi S, Yu L, Tian F, et al. Meta-analysis reveals different functional characteristics of human gut Bifidobacteria associated with habitual diet. Food Res Int. 2023;170:112981.

75. Dziewiecka H, Buttar HS, Kasperska A, Ostapiuk–Karolczuk J, Domagalska M, Cichoń J, et al. Physical activity induced alterations of gut microbiota in humans: a systematic review. BMC Sports Sci Med Rehabil. 2022;14(1):122.

